# Strength of purifying selection on the amino-acid sequence is associated with the amount of non-additive variance in gene expression

**DOI:** 10.1101/2022.02.11.480164

**Authors:** Margarita Takou, Daniel J. Balick, Kim A. Steige, Josselin Clo, Hannes Dittberner, Ulrike Göbel, Holger Schielzeth, Juliette de Meaux

## Abstract

Contemporary populations are unlikely to respond to natural selection if much of their genetic variance is non-additive. Understanding the evolutionary and genomic factors that drive amounts of non-additive variance in natural populations is therefore of paramount importance. Here, we use a quantitative genetic breeding design to separate the additive from the non-additive components of expression variance in 17,657 gene transcripts of the outcrossing plant *Arabidopsis lyrata*. We partition the expressed genes according to their predominant variance components in a set of half- and full-sib families obtained by crossing individuals from different populations. As expected, a population-genetic simulation model shows that when divergent alleles segregate in the population, our ability to detect non-additive genetic variance is enhanced. Variation in its relative contribution can thus be analyzed and compared across transcribed genes. We find that most of the genetic variance in gene expression represents non-additive variance, especially among long genes or genes involved in epigenetic gene regulation. Genes with the most non-additive variance in our design not only display markedly lower rates of synonymous variation, they have also been exposed to stronger purifying selection compared to genes with high additive variance. Our study demonstrates that both the genomic architecture and the past history of purifying selection impacts the composition of genetic variance in gene expression.

## Introduction

Natural populations harbor substantial amounts of genetic variance. However, complex interactions between alleles prevent all but a fraction of this variance from being available to selection; as a result, not all genetic variance is additively transferred to the next generation (Fisher 1918). The higher this non-additive fraction of genetic variance, the more limited in the potential of traits to respond to future selective pressures.

Because theoretical studies predict that selection will deplete additive genetic variance, selection has long been thought to shape non-additive variance (Turelli 1988). To date, evidence that the amount of non-additive genetic variance is shaped by past selective pressures remains elusive (Mousseau and Roff 1987; Crnokrak and Roff 1995; Barton and Keightley 2002; Hunt et al. 2007; Sztepanacz and Blows 2015; Walsh and Lynch 2018). In *Drosophila serrata*, for example, the amount of non-additive variance in wing traits was not related to the extent of genetic variance in male competitive fitness (Sztepanacz and Blows 2015); in maize, in contrast, two traits linked to maize yield appeared to have high non-additive genetic variance (Yang et al. 2019).

Other processes may also shape the relative amount of non-additive genetic variance present in a population (Wolak and Keller 2014). A recent bottleneck, for example, will increase the proportion of additive genetic variance, because some recessive alleles become more frequent, and variation at interacting loci tends to get lost (Willis and Orr 1993; Wang et al. 1998; Waldmann 2001; William G. Hill et al. 2008; Clo et al. 2020). Population admixture can also lead to increased genetic variance (Baranwal et al. 2012; Chen 2013; Vasseur et al. 2019; Clo et al. 2020; Rehman et al. 2021; Baldauf et al. 2022) by increasing the fraction of heterozygotes and opening new opportunities for inter-locus interactions (William G. Hill et al. 2008; Vasseur et al. 2019; Viana and Garcia 2022; Lynch and Walsh 1998; Hahn and Rieseberg 2017). Moreover, the amount of non-additive variance present in populations may also depend on the genetic architecture of phenotypic variation. As the number of loci controlling variation in a phenotype increases, new allelic variants will likely have more opportunities to interact with other variants of the same or different loci to define a new phenotypic value; this new value deviates from the sum of effects of the individual variants and may generate non-additive genetic variance (Lynch and Walsh 1998). Yet, when the genetic basis involves many loci, additive variance is expected to predominate (Mäki-Tanila and Hill 2014).

Because trait measurements are often time consuming, analyses of additive and non-additive variance in populations tend to focus on a few traits only, limiting the ability to systematically quantify the impact of selection and trait architecture on trait variation. The analysis of transcript expression removes this limitation, allowing us to screen the expression level of thousands of gene simultaneously (Ayroles et al. 2009; Romero et al. 2012; Groen et al. 2020). When carried out on full- and half-sib progenies (Fisher 1918; Lynch and Walsh 1998; Huang and Mackay 2016), transcriptome studies can reveal levels of non-additive genetic variance in expressed genes (Leder et al. 2015). Linking the amount of non-additive variance with the selective pressures operating in natural conditions is challenging (Kingsolver and Diamond 2011; Groen et al. 2020). Quantitative genetic analyses of expression have shown that natural selection decreases the amount of genetic variance that can be found in natural populations; therefore, an absence of association between non-additive genetic variance and co-variance with fitness may be inconclusive (Hodgins-Davis et al. 2015). So far, however, methods of population genomics have not been leveraged to determine how the components of genetic variance in expression were shaped by selection over evolutionary time. Indeed, with the advent of population genomics datasets, sophisticated approaches have been developed to identify genomic regions targeted by recent positive selection or to quantify the strength of purifying selection on variation affecting the amino-acid sequence of transcribed genes (Eyre-Walker and Keightley 2007; Stephan 2016; Kim et al. 2017).

We analyze expression variance in 17,657 genes expressed during the exponential growth phase of the outcrossing plant species *Arabidopsis lyrata*. We quantify the additive and non-additive components of genetic variance in the progenies of individuals sampled from two different populations, following a crossing scheme in which diverged haplotypes can segregate within and among families. We apply a model from quantitative genetics to show that this design enhances our ability to detect non-additive genetic variance. We then classify genes by their relative amount of non-additive genetic variance and assess whether they differ in their history of selection by estimating the distribution of fitness effects in each class of genes. We show that the relative amount of non-additive genetic variance in gene expression depends on the history of purifying selective pressures as well as on the architecture of the genome.

## Results

### Inter-population progenies enhance the ability to detect non-additive variance

How much non-additive variance segregates in half- and full-sib families depends on the frequency of heterozygote genotypes (Boeven et al. 2020; Clo and Opedal 2021). When each parent is randomly chosen from two divergent populations, their progeny forms a pedigree with an increased fraction of heterozygotes and carrying alleles that had more time to diverge. We used the Fisher model to simulate population divergence and quantify how much additive and non-additive variance can be expected in the progeny of an inter-population cross(Clo and Opedal 2021). These simulations revealed that the inter-population crosses showed higher absolute and relative levels of non-additive variance, compared to intra-population crosses (Supplementary Figure S1a). This finding is due to the higher frequency of heterozygote individuals harboring two distinct haplotypes compared to intra-population progenies. The levels of additive and non-additive variance detected in inter-population progenies, however, clearly correlated with the levels expected in progenies from intra-population progenies (Supplementary Figure S1b). The progenies of inter-population crosses therefore reveal natural variation in non-additive variance along the genome, similar to that found in classical intra-population crosses, while improving the ability to detect non-additive variation.

### Predominance of non-additive genetic variance in gene expression

We used a set of 10 half- and full-sib families of the outcrossing plant species *Arabidopsis lyrata* ssp. *petraea*. Families were obtained by crossing individuals from two locally adapted populations in Central and Northern Europe (Mattila et al. 2017; Takou et al. 2021) (Supplementary Table S1). We chose to grow plants under controlled conditions reflecting the average temperature experienced in natural environments. We collected leaf material during the exponential phase of vegetative growth and sequenced the transcriptome to quantify gene expression. Half- and full-sib relationships allowed us to partition the total phenotypic variance (VP) of gene expression into additive (VA), non-additive (VNA), maternal (VM), and residual (VR) components of variance in the expression for each transcript in the genome (Figure 1a, Supplementary Table S1). To estimate variance components, we used a Bayesian version of the so-called animal model that fitted additive (relatedness) and dominance (kinship) matrices as well as the identity of the mother plant. In this model, the non-additive component of genetic variation (VNA) comprises genetic variance due to allelic interactions both within and between loci (Caballero 2020). Indeed, the statistical variance due to intra- and inter-locus interactions, i.e. dominance and epistatic variance, cannot easily be disentangled in pedigrees without assuming specific molecular models (Huang and Mackay 2016; Vitezica et al. 2018). We tested whether VNA contributes significantly to genetic variance by comparing the fit of models with and without the dominance matrix for each gene. Based on the Bayesian information criterion (BIC), the incorporation of dominance relationships improved the model for all genes but 22. In comparison, incorporating the maternal parent decreased the BIC for 38.5% of the tested genes (Supplementary Figure S2). In the following, we report on the amount of phenotypic variance (VP) that is explained by the different components of variance in the full model. To facilitate cross-gene comparison, we present components of variance -- VA, VNA, VM, and VR (Supplementary Figure S2) -- as absolute values of each component relative to the total phenotypic variance VP.

**Figure 1:**
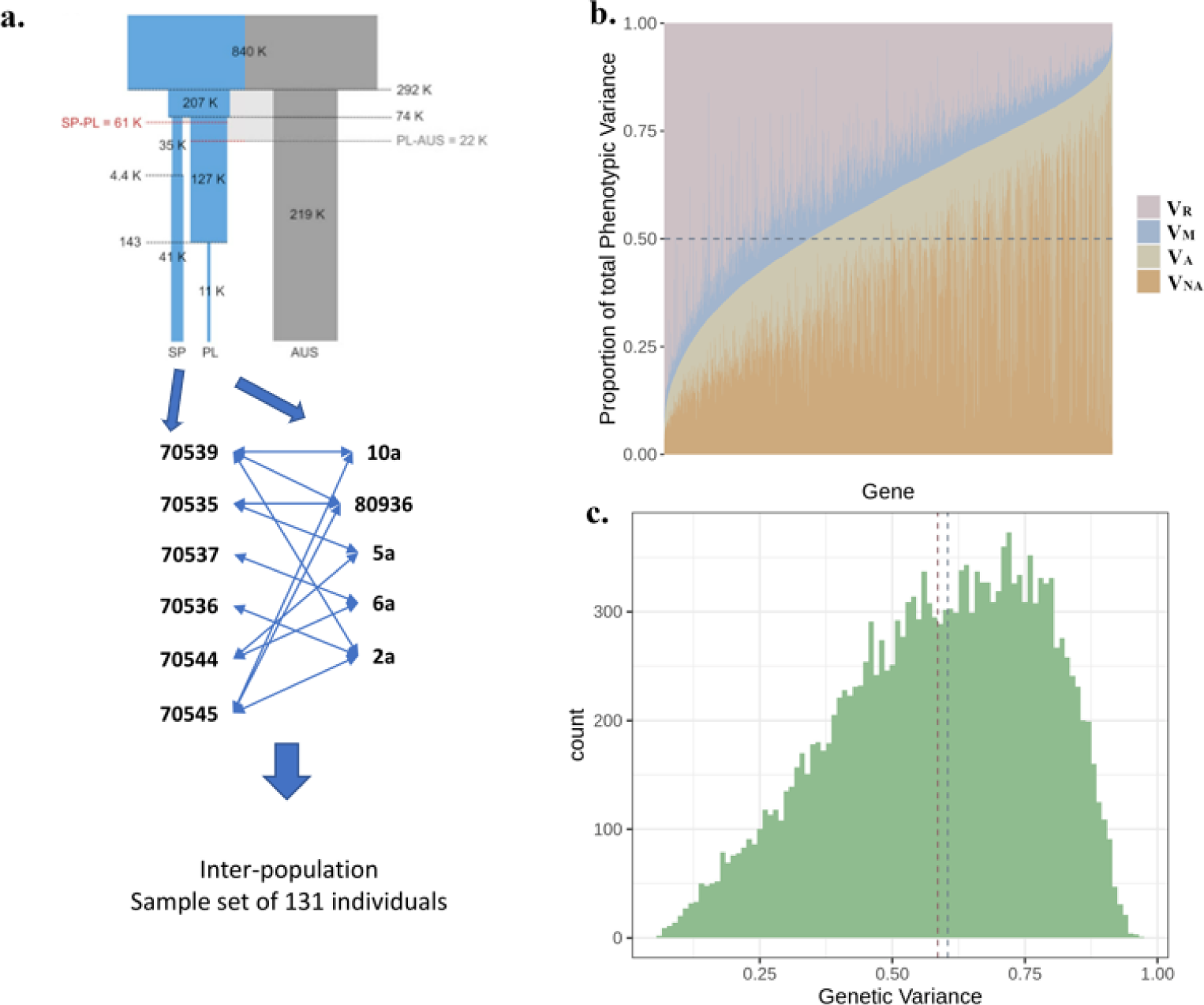
Phenotypic variance of gene expression is mostly composed of non-additive genetic variance. **a.** We took advantage of a set of 131 individuals obtained via crosses between two populations of *A. lyrata*. A previous study had shown that the two populations diverged approximately 61 kyears ago, with the SP populations being more severely bottlenecked than the PL population (redrawn from (Takou et al. 2021a), horizontal lines highlight divergence times). In total, 5 individuals sampled in Plech (PL) were reciprocally crossed with 6 individuals from Spiterstulen (SP). The crosses were done in such a way that plants were reciprocally crossed with 2 individuals from the other population in order to produce full- and half-sib families. **b.** The total phenotypic variance of each one of the 17,657 genes included in the analysis was partitioned into non-additive (VNA), additive (VA), maternal (VM), and residual components of variance (VR). The genes were ordered according to the increasing values of the fraction of total phenotypic variance in the transcript level that was attributed to genetic variance (VG). The dashed line marks the 0.5 point where half of the variance is explained. **c.** The distribution of total genetic variance for each transcript (VG), as a fraction of the total observed variance in expression level VP. The dashed blue line shows the median value (0.604) of the distribution; the dashed red line indicates the mean value (0.585).

In 67.7% of the 17,657 genes analyzed, more than 50% of total expression variance (VP) was explained by the significant contribution of the two genetic components of variance, VNA (mean fraction of VP = 0.375) and/or VA (mean fraction of VP = 0.215). The maternal variance (VM) was close to zero for most transcripts (mean = 0.088). Finally, the origin of the maternal plant did not impact gene expression (all *p* > 2.486e-06, the Bonferroni-adjusted significance threshold). Delineating overall genetic variance showed that VNA contributed the most (25.4% of 17,657 genes had more than 50% of gene expression variance VP due to VNA), in contrast to the additive genetic component VA (6.3% of these genes had more than 50% of their variance in expression explained by VA, Figure 1b-c). Hence, gene expression variance VP is largely due to genetics, but only a limited fraction of that genetic variance is inherited additively in our set of half- and full-sib families. In a control dataset generated by permuting individuals across families, VR explained more than half of the VP for more than 73% of transcripts (mean = 0.717), and VNA was not significant (mean = 0.135), confirming that the levels of genetic variance we report are not random and that, despite the relatively small number of individuals, we are able to quantify non-additive variance in this system.

### Strength of purifying selection associated with the mode of inheritance of regulatory variation

We extended the theoretical expectation of Clo & Opedal (2021) that selection depletes relative amounts of VA and VNA for polygenic traits in a panmictic population, and found that this remains true for VA and VNA measured in inter-population crosses (Figure S1a-c). In addition, the model confirmed that for both intra- and inter-population crosses, the higher the strength of purifying selection, the more VNA contributes to VG at the expense of VA. Selection thus depleted VA more than VNA, and this depletion is detectable in inter-population crosses as well (Figure S1a). These differences are most pronounced for inter-population crosses (non-additive variance explains 8-15% and 33-48% of the genetic variance, respectively, in intra- and inter-population crosses), increasing the statistical power for identifying the effect of selection on components of genetic variance (Figure S1a). Thus, while non-additive effects may appear larger in inter-population crosses than in intra-populations, the effect of selection is qualitatively comparable.

Having determined levels of non-additive genetic variance in gene expression in our dataset, we then used a population genetics approach to determine whether how gene expression is inherited confirms these predictions. We first examined rates at which amino acids evolve in the genes that were expressed in our dataset. We partitioned expressed genes in three non-overlapping sets based on the relative contribution of VA, VNA, or VR to VP, henceforth called high VA, high VNA, or high VR genes (Figure 2a). As a control, we included a fourth set of 3,500 genes sampled at random (control genes) but also intermediate VG (iVG)genes, which did not belong to the high-VA, high-VNA, or high-VR genes. We then estimated the rate at which amino acids evolved between *A. lyrata* and *A. thaliana* across these four groups of genes with different inheritance. The four sets of transcripts differed significantly in their rate of amino-acid divergence. High-VR genes had the slowest rate of amino-acid divergence, whereas high VA genes had the fastest rate (Figure 2b-c). Interestingly, high-VNA genes displayed a rate of amino-acid divergence that was intermediate between those shown by high-VA and high-VR genes.

**Figure 2:**
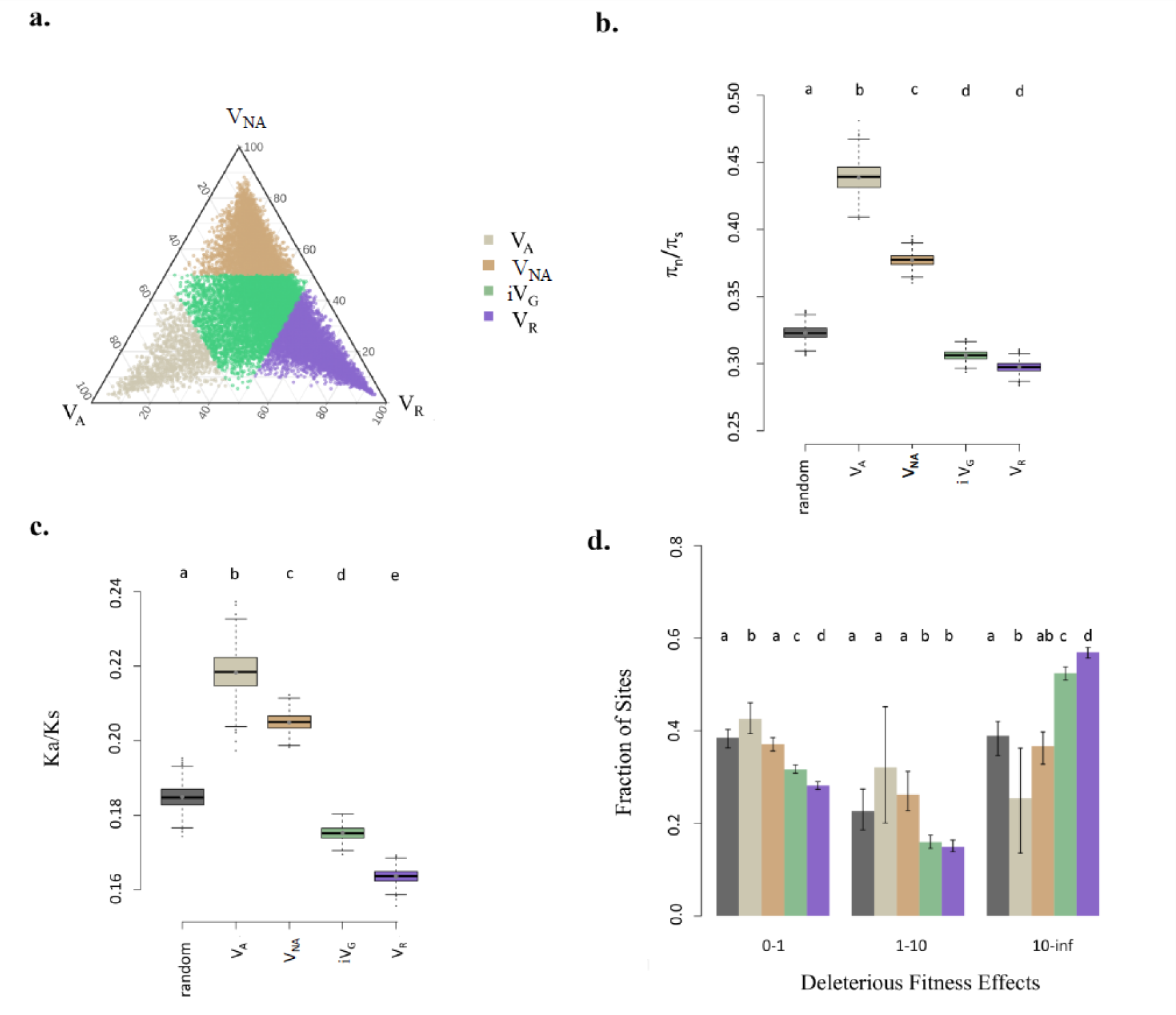
The strength of purifying selection at the amino-acid level is associated with the predominant component of the variance in expression. **a.** We grouped genes based on the partitioning of total phenotypic variance at the transcript level into non-additive, additive, or residual variance. The threshold for each case is 50% of the total phenotypic variance. **b.** We computed the mean and variance of average πi/πs values for each group based on population genomics datasets for PL. For comparison, we added a group of 3500 genes randomly sampled in the genome (gray). **c.** The Ka/Ks ratio for the different gene groups. The confidence intervals and *p* values for b through d were estimated based on 1000 bootstrap replicates. **d.** Inferred distribution of fitness effects of new mutations in the five gene groups for each of three classes of deleterious fitness effect Nes (nearly neutral, Nes<1, weakly deleterious 1<Nes<10, strongly deleterious Nes>10), presented as the fraction of sites hosting mutations that fall in each fitness effect class. **b-d** Mean and variance were computed by bootstrapping 200 times over genes. Black: control group of 3,500 random genes; light gray: genes with high VA contribution relative to VP; orange: genes with high VNA contribution relative to VP; purple: genes with high VR contribution relative to VP; green: intermediate levels of total genetic variance (iVG genes with intermediate values of VA, VNA, and VR relative to VP). Within each plot, the letters indicate whether the values differ significantly from each other.

The rate of amino-acid divergence is expected to increase if positive selection is acting on the encoded protein, but increases may also occur if the evolutionary constraints on the amino-acid sequence are weak. In order to differentiate these alternatives, we used the segregation of molecular variation within species to infer the fitness effect of new mutations and quantify the strength of the evolutionary constraints imposed on the amino-acid sequence. We took advantage of a previous study that determined the demographic history of the same populations of *A. lyrata* (He et al. 2021; Takou et al. 2021). We inferred the distribution of fitness effects (DFE) of new non-synonymous variants in these gene sets following a method described in Takou et al. (2021) that incorporates the specific demographic parameters of each population (Supplementary Figure S3). This method relies on the contrast between the frequency spectrum of synonymous mutations, which are assumed to be neutral, and the frequency spectrum of non-synonymous mutations, which can differ in their fitness effects and have accumulated over long evolutionary time scales (Eyre-Walker and Keightley 2007; Kim et al. 2017). Both the inferred demographic model and the DFE fitted the site-frequency spectra (SFS) of synonymous and non-synonymous variants well (Supplementary Figure S3a), indicating that the model provides a good representation of the evolutionary and selective history of these populations. We present the results for selective estimates obtained for parental population PL; estimates based on variation in SP gave similar results.

These results showed that the inferred strength of evolutionary constraint (Figure 2d; Supplementary Table S2) varies clearly with differences in the type of genetic variance that predominantly explains variation in the expression level of each transcribed gene. Compared to the other sets of genes, the set of genes with low contribution of genetic variance to VP (i.e., high-VR genes) showed a significantly lower proportion of nearly neutral non-synonymous mutations (0 < Nancs < 1, *p* = 0.01) as well as a significantly higher proportion of genetic variants under strong purifying selection (Nancs > 10, *p =* 0.01). This observation indicates that genes evolving under strong constraints at the amino-acid level also tended to show low levels of genetic variance at the expression level.

Our analysis further revealed that selective constraints on high-VA genes are lower than on high-VNA genes (Figure 2c, Supplementary Figure S3b-c). The DFE analysis showed that, compared to other gene sets, high-VA genes hosted a larger fraction of new non-synonymous mutations inferred to be nearly neutral (0 < Nancs < 1, *p*= 0.03). At the same time, a smaller fraction of new mutations was under strong purifying selection in these genes (Nancs > 10, *p* = 0.01). Furthermore, high-VA genes were less likely to map to genomic regions harboring footprints of selective sweeps. Takou et al. (2021) had identified regions in the genome that had both a high probability for a sweep, based on a composite likelihood ratio test integrating the demographic history of the population, and a high F*ST* between the populations. Regions with signatures of selective sweeps extended over 27% of the genome (Takou et al. 2021), but only 17% of the high-VA genes were located in these regions (pairwise t-test, *p*= 2.115e-12), further confirming that these genes are unlikely to contain variants that have been under recent positive selection. Therefore, both the high rate of amino-acid substitutions observed in these genes and their inferred DFE indicate that genes with high levels of additive variance, relative to VP in gene expression, tend to be exposed to weaker purifying selection than are high-VNA genes.

Interestingly, high-VA genes also tend to show a higher mutation rate compared to other genes (Figure 3; Supplemental Figure S3). VA is positively correlated with the rate of nucleotide divergence at synonymous sites, Ks, which reflects the mutation rate per site (ρ = 0.108, p<2.2e-16; Figure 3a-b). VA was also inversely correlated with the rate of genetic divergence between the two parental populations dxy (ρ = 0.1152, *p* < 22e-16; Figure 3a, Table 1).

**Figure 3:**
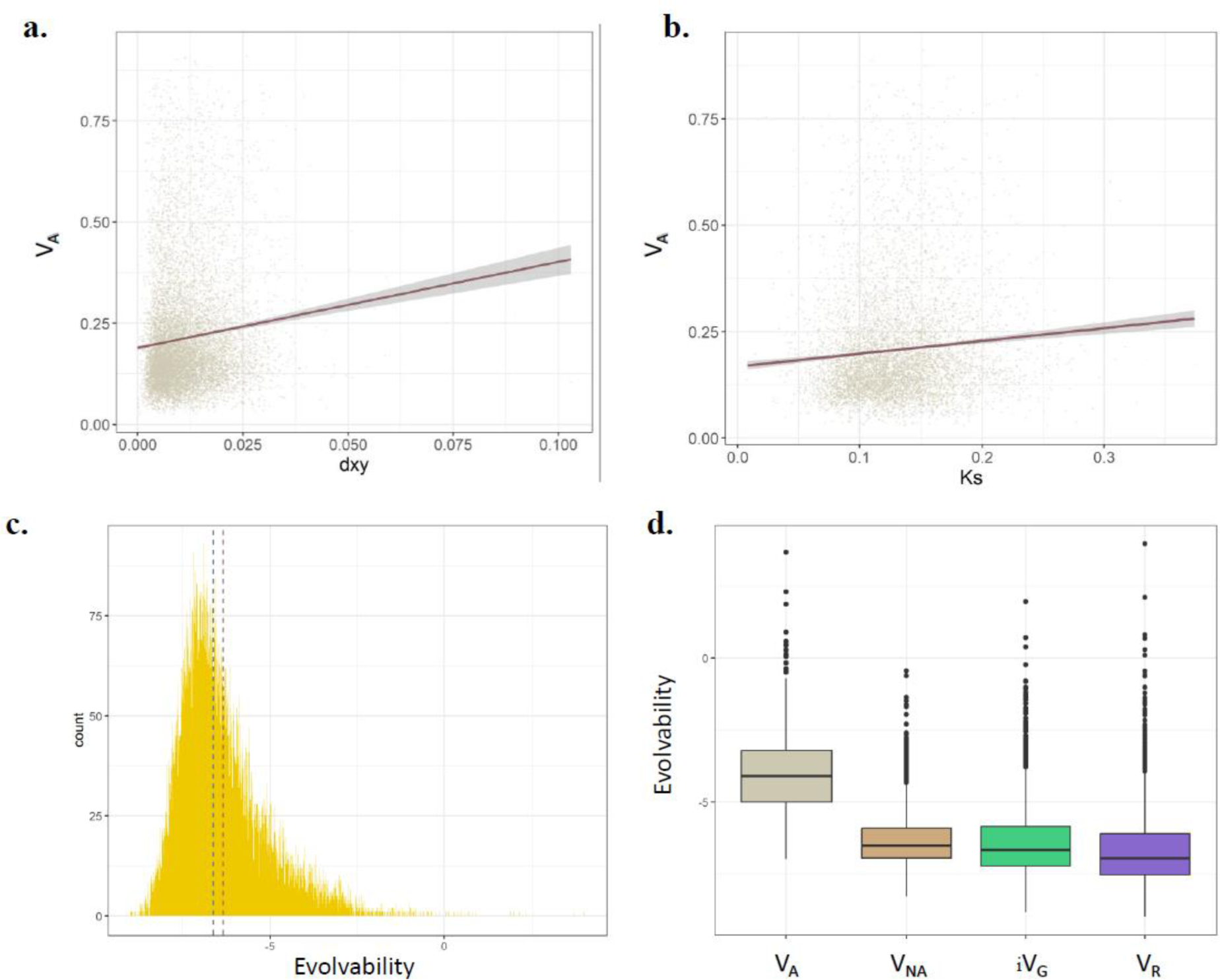
Genes with high VA tend to show higher mutation rate. **a**. Additive variance correlates significantly with the dxy between the genomes of parental populations SP and PL (ρ = 0.1152, *p* < 2.2e-16). **b.** Additive variance is significantly correlated with the per-gene Ks (ρ = 0.108, p < 2.2e-16). **c.** Distribution of the logarithmic evolvability values for each gene. Evolvability was estimated as the unstandardized VA values divided by the logarithmic squared mean gene expression. The red line represents the mean logarithmic evolvability (-6.347) and the blue line, the median logarithmic evolvability (-6.627). **d.** The evolvability for the genes with high VA, high VNA, iVG, and high VR relative to VP. The four distributions are all significantly different.

**Table 1:**
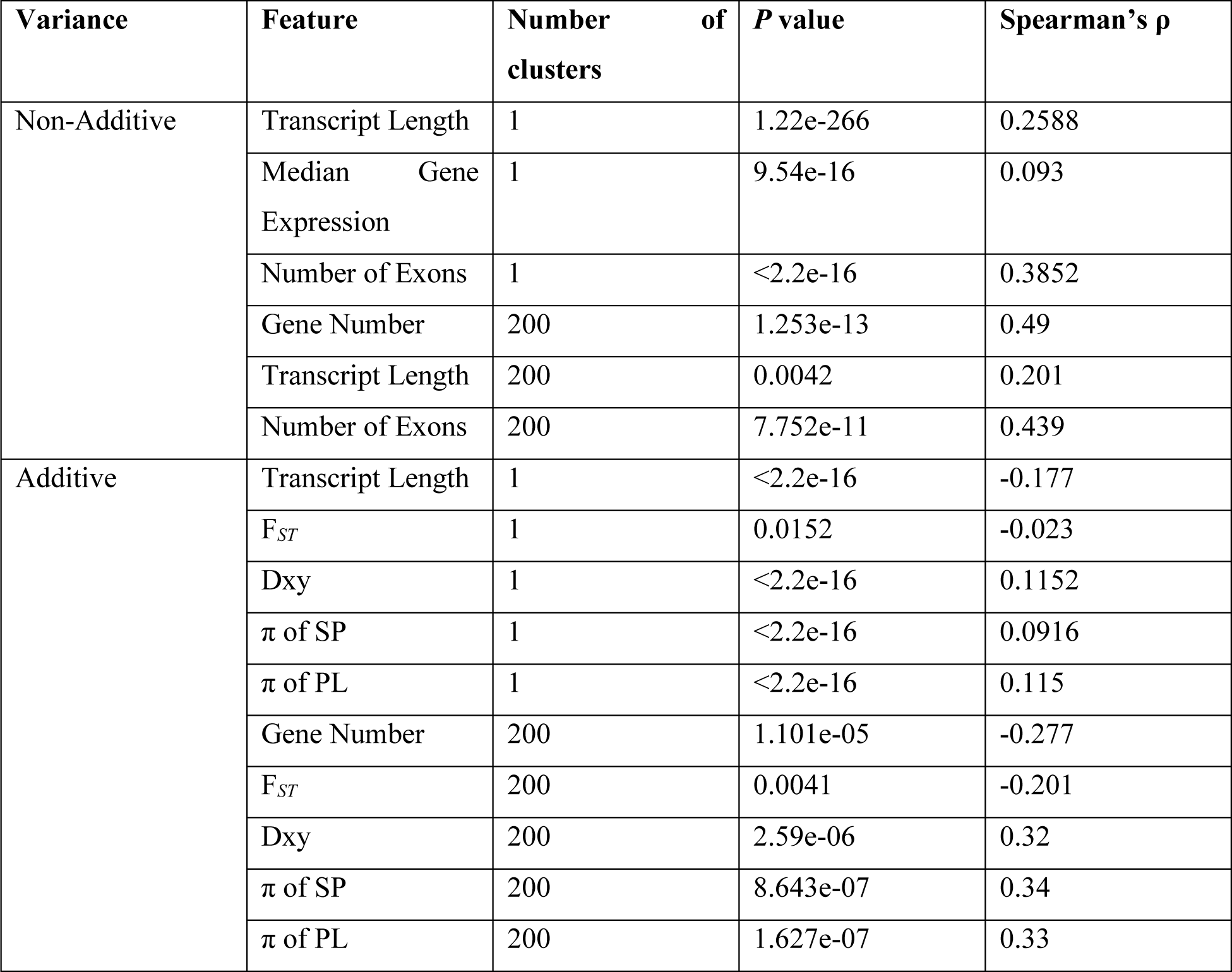
Correlation of dominance and additive variance with genome architecture traits and summary statistics of population genetics. For each correlation between the fraction of variance and the genomic feature, the p value, Spearman’s ρ as well as the number of cluster genes that were grouped, are given. When the number of clusters is 1, the expression level of each gene is treated as independent observation. If it is 200, the clustering algorithm established 200 clusters of co-expressed genes, each with different numbers of genes. In the latter case, correlations were performed on the average of the genes in each cluster.

The evolutionary constraints detected on the amino-acid sequence of high-VNA genes were higher than for high-VA genes but weaker than for high-VR genes, indicating that non-additive genetic variance contributed most to expression variance in genes exposed to intermediate purifying selection pressure. On average, the constraints detected in high-VNA genes were comparable to those experienced in a random set of genes in the genome (Fig. 2c, Supplementary Figure S3d). Interestingly, Ks was markedly lower in genes with high-VNA genes, compared to all other gene groups, and absolute VNA is significantly negatively correlated with Ks (*ρ* = -0.182, *p*<2.2e-16).

### Strength of purifying selection associated with evolvability of regulatory variation

We used the amount of genetic variance relative to the total phenotypic variance to compare the composition of genetic variance across transcribed genes in the genome. This fraction reflects the capacity of a population to respond to selection, but not the magnitude of potential change (Hansen et al. 2011). To estimate the latter, we computed the per-gene evolvability (eμ), dividing the absolute estimate of VA by the logarithm of the squared mean gene expression count (Figure 3c) (Hansen and Houle 2008). Genes that have high relative VA (i.e. Va/Vp) also have significantly higher evolvability eμ (median 0.0165) compared to other sets of genes (all pairwise ks tests, *p<2.2e-16;* Figure 3d). Evolvability eμ increased both with Ka and with the Ka/Ks ratio of the transcript (Spearman correlation coefficient ρ=0.313, *p*<2.2e-6 and ρ=0.276, p<2.2e-16, respectively).

Evolvability eμ was lowest in high-VR genes. High-VNA genes had a significantly lower eμ median (0.00147) than high-VA genes (median 0.0165), but a higher eμ than those with intermediate genetic composition (iVG median 0.0012) or with high VR (0.0009; all pairwise Kolmogorov-Smirnov tests, *p*<2.2e-16, Figure 3d). Therefore, the level of constraints and the mode of inheritance of phenotypic variation modulated both the fraction of VP explained by additive genetic variance and the magnitude of potential evolutionary change in the same way.

### Gene architecture associated with the fraction of non-additive variance in transcripts

In order to better understand whether genomic factors modulate the relative fractions of VA or VNA in VP, we compiled a total of 200 descriptors of genetic variation at each gene; these included the number of known transcription-factor (TF) binding sites in the upstream non-coding region, the length of the genes, and various descriptors of nucleotide diversity within and between the parental populations. We trained a random forest model to identify the genomic architecture or population genetic parameters that best explain variation in the fraction of VA or VNA in gene expression variance and tested the significance of their measured importance by permuting gene identity. We first performed the analysis using the 200 descriptors to predict variation in VA, but the model’s ability to classify high-VA genes was poor (R^2^ = 0.009 and mean square error of 0.021).

For variation in VNA - the fraction of VP explained by non-additive genetic variance-however, the model provided a good classification (R^2^ = 0.18 and mean square error of 0.026), suggesting that descriptors of gene structure are an important factor for classifying genes according to their relative level of VNA (Figure 4a; Supplementary Table S3). Specifically, the number of exons per gene (importance: 53.021; *p* = 0.0164), total gene length (importance: 26.92; *p* = 0.0164), and transcript length (importance: 14.27; *p* = 0.0164) were the three factors that best predicted levels of non-additive variance. The importance of exon number persisted even when gene length was corrected for. Transcript length was significantly correlated with the fraction of VNA (Spearman coefficient ρ = 0.258, *p* = 1.22e-266, Table 1; Figure 4b). We used a linear model to confirm that the effect of transcript length on the relative amount of VNA (*p* < 2.2e-16, see methods) remained significant even after correcting for median gene expression (*p* = 0.0176) and VG (*p* < 2.2e-16). The model did not detect that transcript length and median gene expression interacted significantly (*p* = 0.224). The number of exons per gene correlated significantly with the level of non-additive variance (ρ = 0.358, *p <* 2.2e-16; Figure 4c). The random forest analysis further revealed that the density of SNPs within a gene was a predictor of high-VNA. Strikingly, all significant descriptors tended to increase the mutational target size of the transcript. We thus conclude that an increased fraction of VP explained by non-additive variance is likely to be observed as the mutational target size increases.

**Figure 4:**
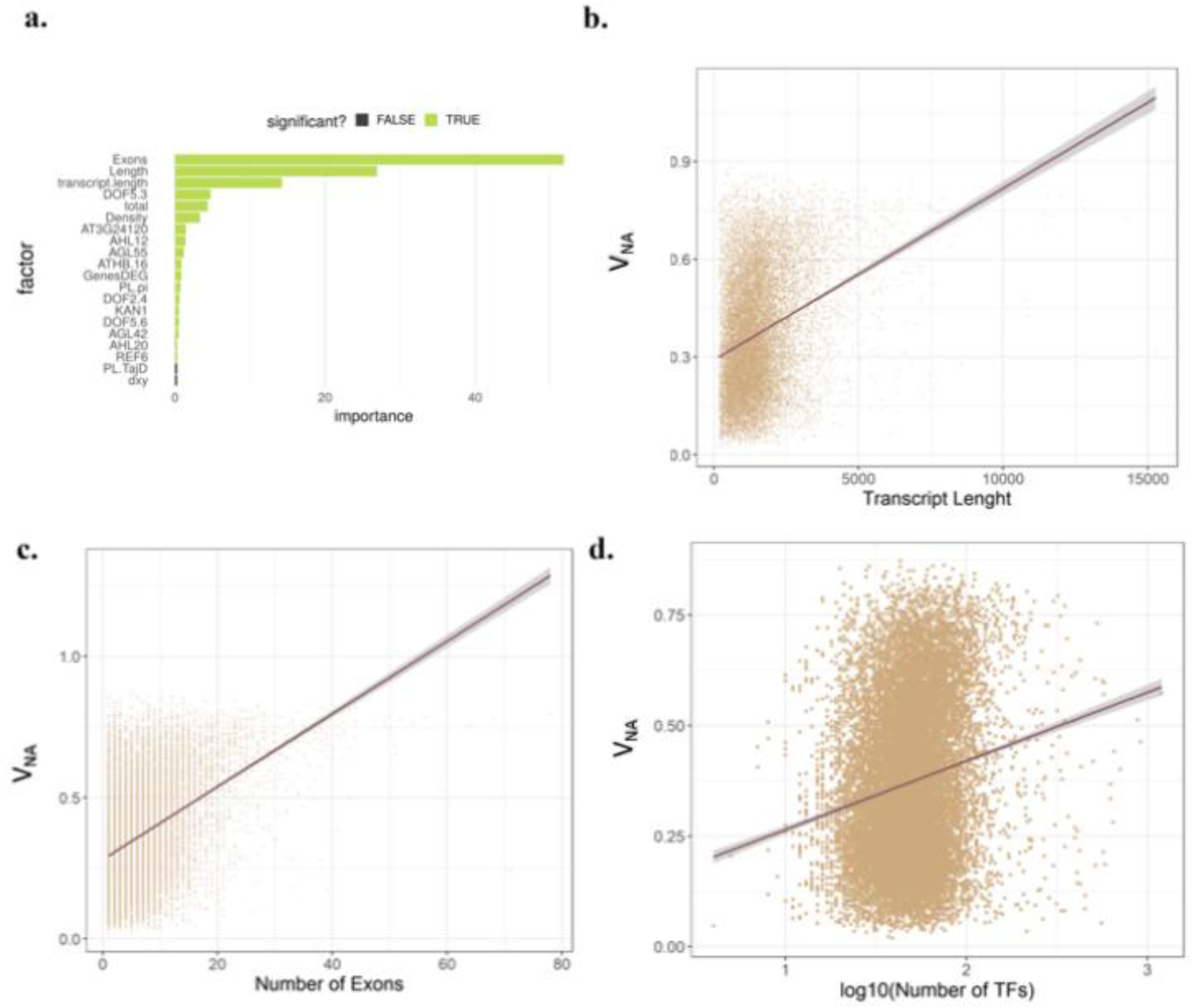
The level of non-additive variance depends on gene architecture. **a.** Importance of tested genomic parameters on the level of non-additive variance each gene. The twenty factors that explain the most variance are shown. The predictive power of the model is R^2^ = 0.18. Significant parameters are shown in green. **b.** The fraction of non-additive variance correlates significantly with the transcript length of each gene (*p*=1.22e-266, ρ=0.258). **c.** The fraction of non-additive variance correlates significantly with the number of exons that a gene has (*p* < 2.2e-16, ρ=0.385). **d.** The fraction of non-additive variance correlates significantly with the total number of transcription-factor binding sites detected in the upstream vicinity of the gene (*p* < 2.2e-16, ρ=0.162).

Interestingly, analysis further revealed that the total number of TF binding sites explained relative VNA well (*p* = 0.0163; importance 4.34; Figure 4d). In particular, the transcription-factor binding sites that are most significantly associated with high-VNA values were putative targets of a DOF zinc finger protein expressed in young rosettes (DOF 5.3; importance 4.722; *p* = 0.0163); this protein (AT5G60200 TMO6) was recently shown to drive the stem cell division that is necessary for plants to thicken (Miyashima et al. 2019). This regulator was likely active in the tissue samples we collected, because we sampled plant material during the exponential phase of rosette growth.

We further quantified the distribution of genes with high VNA across gene ontology (GO) categories of expressed genes and tested for specific functional enrichment by permutation (see methods). We found that high-VNA genes were enriched for GO categories reflecting cell differentiation. The top enriched GO terms for the highest estimates of relative VNA were “chromatin silencing” (*p* = 4.5e-16) and “cytokinesis by cell plate formation” (*p* = 2e-11), “DNA methylation” (*p* = 4.3e-11; Supplementary Table S4). In addition, genes with high VA were not randomly distributed among molecular functions; unlike genes with high VNA, the functional enrichments were less striking and not related to cell differentiation. Among the top GO categories, the most enriched among genes with high VA were several functions linked to central metabolism such as “glucose catabolic process” (*p* = 8.3-05) and “glycolytic process” (*p* = 1.6e-05), but we also detected enrichments among genes involved in the “response to cadmium ion” (*p* = 0.00015) and “proteasome core complex assembly” (*p* = 0.00016; Supplementary Table S5).

### Non-additive variance relative to VP increases with the size of the co-regulatory network

Because we found that VNA (relative to Vp, as above) was associated with the putative activity of a transcription factor, we investigated whether VNA depended on the architecture of the network of gene co-regulation. We computed clusters of co-regulation. As expected for a transcriptome dataset, many genes formed co-expressed clusters of variable size (Supplementary Figure S2). Yet, major genetic incompatibilities or large-effect variants were unlikely to explain the data (Bomblies and Weigel 2007). Indeed, a genome-wide association study (GWAS) for each expressed gene showed that most of the genetic variation in expression could not be traced to large-effect loci, as significant associations were detected for only 174 out of 17,657 tested genes (Supplementary Figure S2). Therefore, phenotypic aberrations and broad regulatory alterations sometimes seen in inter-cross populations due to genetic incompatibilities are unlikely to explain the data (Alcázar et al. 2010; Bomblies et al. 2010; Lafon-Placette and Köhler 2015; Todesco et al. 2016).

We further analyzed how the composition of genetic inheritance varied across clusters of co-expressed genes. We found that the median VNA in a cluster of co-regulated genes increased with the number of genes contained in this cluster (ρ = 0.49, *p* = 1.253e-13, Supplementary Table S6); additionally, the correlation between the gene number per cluster and median VA was negative (ρ = -0.277, *p* = 7.101e-05).

Taking co-expression into account did not alter the results described above (Table 1). The median VNA per cluster was positively correlated with the median transcript length (ρ = 0.201, *p* = 0.0042) and median gene exon number (ρ = 0.439, *p* = 7.752e-11). The median VA of each cluster was negatively correlated with the median value of relative population divergence F*ST* (ρ = -0.201, *p* = 0.0041, Table 1) and positively correlated with the absolute measure of population divergence dxy (ρ = 0.320, *p* = 2.59e-06) among parental populations. The median VA was also correlated with the median π values in both populations (SP: ρ = 0.34, *p* = 8.643e-07, PL: ρ = 0.33, *p* = 1.627e-07), and high-VA genes were among the genes that displayed the highest divergence at both synonymous and non-synonymous sites (Supplementary Figure S3b-c). Accordingly, our data suggest that both inter-allelic and inter-genic interactions increase the amount of non-additive variance in gene expression; this result is expected given that this variance includes both dominance and epistatic genetic variance.

## Discussion

Delineating the genomic and evolutionary factors that increase the non-additive components of genetic variance is essential for understanding the genetic architecture of natural variation and how it might fuel future adaptation. Non-additive variance, a statistical component of genetic variance, contributes to similarity between siblings, but trait variation based on non-additive variance is not inherited by the siblings’ offspring and thus cannot help the population respond to selection (Fisher 1918). Estimating levels of non-additive variance is notoriously difficult, because it requires both large sample sizes and the ability to control various sources of phenotypic variance (Sztepanacz and Blows 2015; Hivert et al. 2021). Here, we used a quantitative genetics model based on differential kin relationships, an approach argued to be the most accurate for quantifying non-additive variance (Huang and Mackay 2016). Plants are especially suited for dissecting genetic variance because both genetic crosses and environmental variance can be completely controlled (Lynch and Walsh 1998; Yang et al. 2019). In neutrally evolving traits, levels of non-additive variance are predicted to be low compared to additive genetic variance (William G. Hill et al. 2008; Hivert et al. 2021). Expanding the model proposed by Clo & Opedal (2021), we can show that non-additive variance will be more prevalent in inter-population families than in families obtained by panmictic crosses within populations. At the same time, levels of non-additive variance detected with intra- and inter-population progenies are expected to correlate. Using inter-population progenies thus improves our ability to quantify genomic variation in non-additive genetic variance and facilitates the study of its genomic and evolutionary drivers.

The estimates of non-additive variance that we report for 17,657 gene transcripts, reveal that non-additive variance forms a substantial fraction of the regulatory variation quantified in our study. Clearly, gene expression levels are subject to natural selection (Ayroles et al. 2009; Hodgins-Davis et al. 2015; Kremling et al. 2018; Groen et al. 2020). Linking trait variance with the history of selection these traits have experienced is a difficult task (Lynch and Walsh, 2018). Here, we address this challenge by computing the distribution of fitness effects (DFE) of deleterious mutations in groups of genes that differ in mode of inheritance of gene expression (Eyre-Walker and Keightley 2007; Takou et al. 2021). Our data show compellingly that purifying selective forces that have operated over long evolutionary time scales in the past impact the levels of non-additive variance measured with an inter-population pedigree.

Genes whose genetic variance at the transcript level is predominantly non-additive in inter-population progenies were significantly more constrained than genes showing a large fraction of additive variance. The most strongly constrained genes were found among the genes that were almost completely depleted of any genetic variance in transcript level, additive and non-additive variance alike. Both levels of constraints measured within populations and between species confirmed this pattern. It was also maintained regardless of whether genetic components were standardized to the gene expression variance VP or to the log of the gene expression mean. Although levels of genetic variation in expression have long been known to be low in genes with constrained amino-acid evolution (Fay and Wittkopp 2008; Ayroles et al. 2009; Romero et al. 2012; Hodgins-Davis et al. 2015; Hämälä et al. 2019), as far as we know, we are the first to report that non-additive transcript variance increases with the strength of purifying selection but ultimately decreases in genes exposed to the strongest stabilizing selective forces. These differences are revealed by a skew in the frequency spectrum of non-synonymous mutations in both parental populations as well as in the number of amino-acid substitutions that ultimately fix in a population. Indeed, selection against recessive deleterious variants will tend to maintain them at low frequency, in the realm where non-additive variance can make for a larger fraction of genetic variance (Caballero 2020).

Our data reveal an important additional feature: the rate of synonymous divergence and polymorphism is significantly lower in genes with high non-additive variance compared to all other groups of genes, suggesting that variation in the mutation rate may also affect the composition of genetic variance for expression. Additionally, the genes with a high relative amount of VA, which were also those with fewer constraints on the amino acid sequence and possibly higher mutation rates, showed higher evolvability than the genes with a high fraction of non-additive variance. This shows that the evolutionary potential, which is highest in these genes, decreases as non-additive variance explains more of the phenotypic variance (Hansen and Houle 2008; Hansen et al. 2011).

We further identify genomic features that favor non-additive variation in gene expression. Non-additive variance in gene expression was promoted by transcript length, suggesting that the mutational target size of the expressed locus favors the emergence of such variance. Our study of transcript-level variance further indicates that non-additive variance increases as the network of co-regulated genes increases, which confirms previous observations that proteins that are more central tend to be more constrained at the amino-acid level (Hahn and Kern 2005; Orr 2009; Josephs et al. 2017; Masalia et al. 2017). In addition, this work shows that what promotes non-additive genetic variance is distinct from the molecular interaction between alleles. Indeed, contrary to our findings, additive molecular effects on fitness (e.g. co-dominant alleles) were inferred to be more frequent in highly connected genes of the same species (Huber et al. 2018).

Finally, we report that, in the inter-population progenies used here, non-additive variance is particularly important among genes influencing the epigenetic remodeling of chromatin in differentiating cells. We associated non-additive variance in expression with the presence in the promoter of DNA motifs bound by a regulator of stem thickening (Lister et al. 2004). Although epigenetic variation has been often hypothesized to facilitate adaptation (Schmid et al. 2018), our results indicate that given how the expression variance of epigenetic regulators is regulated, these regulators are potentially less likely to produce additive variance than other molecular components of the genome.

In summary, this study identifies clear genomic and evolutionary drivers of non-additive genetic variance in gene expression. We report that, on average, non-additive variance explains 37% of the genetic variance in gene expression. These conclusions, however, rely on inter-population pedigrees: compared to intra-population pedigrees, inter-population pedigrees are predicted to increase total levels of genetic variance in gene expression. Accordingly, although non-additive genetic variation might be somewhat overestimated in our population cross, non-additive genetic variation is larger and more important than commonly assumed. Indeed, a meta-analysis concluded that the fraction of non-additive variance that had panmictic intra-population pedigrees reached an average of 38% of the genetic variance, a value close to the value we report here, and higher than expected (William G Hill et al. 2008; Wolak and Keller 2014). A previous study, using sticklebacks, determined that non-additive genetic variance also formed about 50% of the total genetic variance in transcript levels (Leder et al. 2015). For life-history traits in non-domesticated species, where allelic frequencies are often skewed by the history of bottlenecks and demography, this fraction reached 60% of the genetic variance (Wolak and Keller 2014).

This large-scale transcriptome study is, to our knowledge, the first to associate estimates of purifying selection with the quantitative genetics of gene expression. Results show that quantitative genetics, which aims to predict short-term responses to selection, can be integrated with population genetics, a field focused on reconstructing historical evolutionary processes. Indeed, we report that past stabilizing selection within populations is associated with an increased contribution of non-additive genetic variance to transcript level variation and a decrease in evolvability. This result, however, was obtained by measuring components of inheritance in inter-population progenies. In such progenies, haplotypes that have diverged over time are allowed to segregate within and between families. Our model indicates that this crossing design enhanced our ability to detect non-additive genetic variance as well as its depletion through selection. The model also indicated that levels of non-additive variance in inter-population pedigrees correlate with those expected in natural populations. The fact that the relative amount of non-additive variance we measured correlates with the strength of purifying selection detected within populations is in agreement with this prediction and suggests that levels of non-additive variance present in natural populations will be highest among genes experiencing intermediate selective constraints. Future studies will have to carefully test whether intra-population progenies in this and other plant or animal species exhibit similar patterns. Whether stabilizing selection has eroded the additive genetic variance and evolvability of transcript levels in other tissues, developmental stages, or conditions will also be important to examine.

## Methods

### Plant Material Preparation

Six plants from an *Arabidopsis lyrata* ssp. *petraea* population located in Spiterstulen, Norway (SP; 61.41N, 8.25E) were crossed with two of six plants from Plech, Germany (PL; 49.65N, 11.45E), to obtain inter-population F1 offspring progenies. For crossing, parental individuals were propagated clonally and vernalized for nine weeks at 4°C and 12-hour daylength, and subsequently transferred to the greenhouse (16-hour daylight, 22°C) until they flowered. Each individual from one population was then crossed with two individuals from the other population (Figure 1a). All crosses were reciprocal -- each individual plant was used as both pollen receiver and donor -- to be able to control for maternal effects. Since the same plants were used for the reciprocal crosses, the resulting seeds are considered full-siblings and they are termed “family”. All family members are half-siblings with the members of at least another family that share the same parent (Supplemental Table S1).

Seeds obtained from each reciprocal cross were stratified on wet filter paper in the dark at +4°C and kept under these conditions for 4 (SP seeds) and 7 days (PL seeds). Subsequently, the seeds were allowed to germinate at 20°C, 16 hrs of daylight. Once the cotyledons were fully open, the plants were transferred to pots and placed in a walk-in growth chamber (Dixel, Germany) set at 12°C, 16 hrs of daylength. The light intensity was adjusted via the light ratio of the LEDs (LED Modul III DR-B-W-FR lights by dlicht®). The LEDs were set at 100% intensity of blue (440 nm), red (660 nm), and white light with total measured intensity of 224 +/- 10 μmol * sec^-1^ *m^-2^ . During both experiments, a light pulse of far red (750 nm) was implemented for 10 mins at the end of the day. We sampled the above-ground leaf material of each individual when the 10^th^ leaf was visible on the rosette. This time point was chosen because previous observations had shown that it is the stage at which plants grow exponentially. All sampling was performed at 4 hours Zeitgeber time. Two families (annotated as fam11 and fam12 in Table S1) did not germinate well, and the remaining individuals died before producing the 10^th^ leaf; thus these families were removed from subsequent analysis.

### RNA extractions for transcriptome sequencing

To quantify the relative importance of additive and non-additive variance in gene expression, we extracted RNA and DNA using the AllPrep DNA/RNA Mini Kit (Qiagen, USA). We first used the extracted DNA to confirm that all the samples were crosses between the two populations, by heterozygosity at one diagnostic locus. For confirmation, we used custom primers amplifying DNA around a region that the restriction enzyme PvuII could not digest at a specific position within one of the two populations. The forward primer sequence was 5’- GCACAAGACTGCTGTAACGC-3’ and the reverse primer sequence was 5’- AATGGCCTCCCGTATTTGCA-3’. The amplified region was digested and visualized on a 0.8% agarose gel. A sample from SP or PL would have a set of bands unique to the respective parental population, whereas a hybrid between the populations would show all possible bands. RNA quality and quantity was assessed with Qubit 4 Fluorometer (Thermofisher Scientific, Germany), 2100 Bioanalyzer (Agilent, USA) and a 0.8% agarose gel. At this stage, any RNA extract that showed signs of degradation was discarded.

### Sequencing and transcriptome data preparation

We obtained high-quality RNA samples for 133 individuals. RNA was sequenced on Illumina HiSeq4000 at the Western German Center for Genomics (Cologne, Germany) following the TruSeq protocol. Sequence reads were paired-end, 75bp-long, and stranded. They yielded on average 80X coverage of the transcriptome of each sample. Sequence quality was assessed with FastQC. The first 3 bp and unpaired reads were discarded with Trimmomatic v0.36 (Bolger et al. 2014). Transcriptome mapping against the *A. lyrata* v1 (Hu et al. 2011) reference genome was performed with STAR v2.5.3 (Dobin et al. 2013) using standard settings plus a cut-off for maximum indel length of 10 kbp. The ribosomal RNA was identified and removed based on the *A. lyrata* genome annotation v37 and using bedtools subtract (Quinlan and Hall 2010). We calculated the gene counts per individual using htseq-count (Anders et al. 2015) and estimated the number of fragments per kilobase per million reads (FPKM) in R. Based on the total FPKM and log2 distribution of the gene counts, we removed two individuals with low FPKM count and divergent log2 distributions (Supplemental Figure S1) from the analysis; both showed indications of low transcriptome sequencing quality.

All reciprocal crosses were done in the greenhouse, which is not pollinator free. To rule out unintended crosses, we used variant-calling to assign parents. The single nucleotide polymorphisms (SNPs) of the hybrids’ transcriptomes as well as of the parental genome sequences (Takou et al. 2021) were called using samtools v1.5 and bcftools v1.5 (Li et al. 2009). The sites were filtered based on their quality (minimum 60), depth (minimum 10), and minor allele frequency (minimum 0.01), using vcftools v0.1.15 (Danecek et al. 2011). All indels were removed and no missing sites were allowed. We used custom python scripts to estimate the genetic distance between the hybrids and the parental genomes. The genetic distance between two lines was thus estimated as:

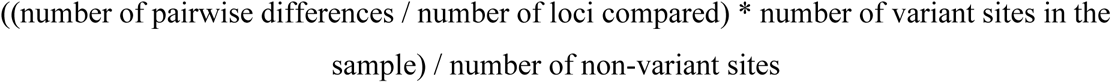

Furthermore, the relatedness between all individuals was estimated with the vcftools build in function. A PCA of the samples was performed with R Statistical Language (R Core Team 2018), using the adegenet package (Jombart 2008). At the end of the analysis, we had to correct the family of origin for 22 samples (Supplemental Figure S6).

### Partitioning of gene expression variance into its components

We partitioned the expression variance of each gene into its genetic and environmental components. To this end, we first filtered genes based on their expression level. Genes with total reads less than 130 across all individuals, genes with median counts of more than 180,000, and genes with FPKM of less than 0.31 in 80% of the individual were removed from the analysis. For each of the remaining genes, an animal model (Lynch and Walsh 1998; Wilson et al. 2010) was fitted in R (R Core Team 2018) using the MCMCglmm (Hadfield 2010) package. MCMCglmm takes a Bayesian approach to fitting general linear models. The fitted models included the population of the mother plant as a fixed effect to account for variation associated with the mother plant’s population of origin. Random effects included the additive and dominance matrix, which was calculated using the nadiv package in R (Wolak 2012). Additionally, the identity of the mother plant was included to control for effects of the plant that the seeds matured on. The model, thus followed the following formula:

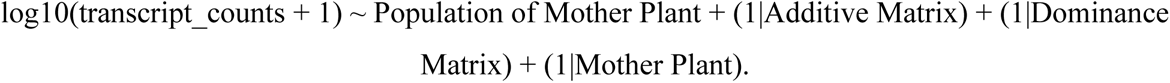

The prior distribution for both fixed and random effects was an inverse gamma (0.001; 0.001). The counts were logarithmically transformed into log10 (counts + 1) to approximate a Gaussian distribution. Each model was run twice for 2.2 million iterations; from these, the first 200,000 iterations were discarded and every 2,000^th^ iteration was sampled. To correct for poor model fit, we removed from the dataset genes that had a posterior effective sample size of less than 500 for at least one of the random or fixed effects, as this indicated that posterior samples were not sufficiently independent. Additionally, we removed from the analysis genes for which the Markov chains had a low quality of convergence, i.e. with Gelman Rubin criterion values greater than 1.1 as recommended in (Gelman and Rubin 1992). In the end, we used the expression counts of 17,657 genes in the downstream analyses.

For each gene, the additive (VA), maternal (VM), non-additive (VNA), and residual (VR) variance components were extracted from the model. The sum of the four variances comprises the total phenotypic variance (VP), whereas the sum of the VA and VNA represents the total genetic variance (VG). To compare the genetic composition of phenotypic variance across the genome, we present VA, VNA, VM, and VR as fractions of VP, unless stated otherwise. We also computed the evolvability (eμ) of each gene as the absolute value of additive variance divided by the log10 of squared mean gene count (Hansen and Houle 2008). The analysis was also implemented for the growth rate of each plant, which was defined as the number of days between potting and the appearance of the 10^th^ true leaf. Furthermore, for each gene we ran two additional linear mixed models; one excluding the dominance matrix from the random effects and one excluding the maternal effects from the random effects. Then, we compared the reduced model to the more complex model described above, and the best-fitting model was selected based on the Bayesian information criterion (BIC). Furthermore, we permuted the membership of the plants in families 100 times, to estimate the variances under random genetic relationships.

### Assessing the impact of inter-population crosses on non-additive genetic variance

In this study, we used families obtained by crossing individuals from different populations. To get a sense of whether population inter-crossing distorts the amount of additive and non-additive variance compared to crosses within a natural population, we simulated the evolution of allelic haplotypes during population divergence. We adapted the model of Clo and Oppendal (2021), which is a derivation of the Fisher geometric model with dominance effects at the phenotypic scale. In each simulation, we modeled two populations (Pop 1 and Pop 2) at mutation-selection-drift equilibria that have evolved separately and accumulated genetic differentiation; other demographic (population size) and genetic parameters (mutation rate U=0.05, mutational effects) were set to be equal. At each locus, each population has maintained an ancestral optimal allele (called A, maintained at frequencies p1 and p2), but they could also accumulate deleterious mutations with potentially different effects on fitness in the two populations (alleles B1 and B2 found at frequencies q1 and q2 respectively). The additive and dominance effects of each mutation were drawn from a Gaussian distribution with mean 0 and variance 0.05 for additive effects and mean 0.25 and variance 0.05 for dominance effects. We then mimicked two crossing designs: progenies obtained from two parents chosen randomly from population P1, and progenies obtained by crossing two parents chosen randomly from P1 and P2.

For the intra-population progeny, 3 genotypes are possible at each locus

GAA with F(AA) = p1²

GAB1 with F(AB1) = 2.p1.q1

GB1B1 with F(B1B1) = q1²

For the inter-population progeny, up to 4 genotypes are possible:

GAA with F(AA) = p2(p1 + q1².q2)

GAB1 with F(AB1) = p2².q1 + (p1q1). (p2q2)

GAB2 with F(AB2) = p1q2

GB1B2 with F(B1B2) = q1q2

It follows then that the fraction of heterozygous individuals is higher in the inter-population progeny. We estimated both absolute and standardized VA and VNA (= VD in this model as there is no inter-locus directional epistasis on either phenotype or fitness) both within populations and in the inter-population crosses. In this case, because VR did not vary, VA and VNA were standardized to total genetic variance VG. We simulated three levels of purifying selection: strong, weak, and neutral (modeled by changing the stabilizing selection parameter ω² = 1, 20, and 100, respectively). A more complete description of the model in available in *(35)*.

### Gene ontology enrichment analyses and clustering of gene expression

Gene ontology enrichment analyses were performed in R with the topGO package (Subramanian et al. 2005; Alexa and Rahnenfuhrer 2016), using as gene universe the list of annotated orthologue genes in the *A. thaliana* genome and applying the test to genes ranked by decreasing values of either VA or VNA. The enrichment was analyzed by Fisher’s exact tests with a *p* values threshold of 0.0381 and 0.038, which was derived by permuting gene identities 1,000 times.

In order to test for potential clustering of the data and therefore non-independence of the expression phenotypes, we first computed the pairwise Spearman correlation coefficient between gene expressions across all the gene pairs in the dataset. To cluster the genes, the pairwise *ρ* values were transformed to pairwise Euclidean distance with the formula D = (1-*ρ*)/2. Hierarchical clustering was done in R with the function hclust (Müllner 2013). Potential clusters were identified by examining the within-group sum of squares for clusters 1 to 300. Clusters were pruned to 23, 50, 100, and 200 clusters by R function cuttree. The impact of clustering on the genetic variances and gene properties (described below) was tested by correlating the median VA, VNA, or transcript length of each group of clustering genes with the number of genes within each cluster.

### Random forest analysis

We asked whether there are relationships between the components of genetic variance and TF binding sites, gene properties, and population genetic features of *A. lyrata* genes. We assembled all molecular features in an sql database. Putative binding sites were predicted from *A. thaliana* using JASPAR2018 Bioconductor package (JASPAR 2018). The generated binding-site sequence matrices were then matched to the aligned regions of *A. thaliana, A. lyrata, A. halleri,* and *Capsela rubella*. We subsequently extracted counts of each transcription factor binding-site in the 1 kb upstream of each gene of *A. lyrata*. We treated the count of each TF binding site per gene as a separate feature and determined the total number of TF binding sites per gene, the number of exons, the length of the transcript and gene. Furthermore, we determined the mean value of the following population genetic features for each gene: F*ST*, Tajima’s D, SNP density per kb, dxy and π (per population of origin) as estimated in (Takou et al. 2021). Additionally, we categorized each gene based on whether it was located within a selective sweep area defined in (Takou et al. 2021) and whether the gene was differentially expressed between at least one pair of families. Differentially expressed genes (DEGs) were identified using the R library DESeq2 (Love et al. 2014), while simultaneously accounting for family effects and using *p* values corrected for multiple testing. Taken together, these descriptors of genomic variation provided a total of 434 features in which we could assess the predictive impact on the composition of variance (Table S3).

We trained a random forest model using the ranger library in R (Wright and Ziegler 2017) with VA or VNA as the response variable and all features of the sql database as predictor variables. We set the number of trees of the forest to 500, the number of variables to possibly split at each node (mtry) to 200, and the impurity mode to impurity_corrected. From the trained random forest, we extracted the relative importance (i.e. explanatory power) of each predictor variable as well as the out-of-bag prediction error, which is informative about the predictive power of the model. Finally, we estimated a *p* value for each predictor variable using the “Altmann” method with 60 permutations.

We further explored the impact of the genome architecture on VA and VNA by correlating estimates of VA, and VNA with the transcript length and gene length, using Spearman’s rank coefficient. We also quantified the combined effect of transcript length and median gene expression in a generalized linear model including the level of broad sense heritability (fraction of VG explaining VP) as a co-variant, and fitted with the GLM (generalized linear models) function and Poisson distribution of the residual variance. Finally, we also tested whether average gene length varied significantly across enriched GO categories with a mixed GLM model, using the lme4 function (Bates et al. 2015) with variance component fraction of VP (VA or VNA) as fixed effect and the category “gene ontology” as a random effect, to correct for pseudoreplication.

### Distribution of fitness effects across gene classes based on predominant genetic variance components

To investigate the level of constraint of the groups of genes with different levels of genetic variance in the PL and SP, the parental populations of our crosses, we grouped the genes in four categories based on whether more than half of VP was due to additive, non-additive or residual variance, grouping all other genes in an intermediate group iVG, with intermediate fractions of all three variance components. A fifth set of 3,500 random genes was further added as a control group. We assessed the distribution of fitness effects in each group of genes (DFEs) using a combination of general python functions, δaδi and fitδaδi (Gutenkunst et al. 2009), following the exact procedure we had previously reported in (Takou et al. 2021), and for which a code is available. In brief, we used synonymous variation to determine the best-fitting demographic parameters to the parental population PL (or SP). We then predicted the distribution of fitness effects given the simplified demographic model in each population and for each group of genes. The simplified demographic model was inferred by maximizing the composite likelihood of the folded SFS at 4-fold degenerate sites of each gene group in PL (or in SP) using the “L-BFGS-B” method and basin-hopping algorithm implemented in scipy (Virtanen et al. 2020). These models provided a good fit of the predicted neutral SFS to the data of each gene set. After estimating the demographic parameters, we proceeded to the second step of our analysis and used the 0-fold SFS of each gene set to fit their respective DFE by estimating the shape and scale parameter of a gamma distribution of selection coefficients. Analysis was performed assuming that deleterious variants were all co-dominant (*h* = 0.5). For this, we estimated the 4-fold population-scaled mutation rate theta, which reached 24,000 for PL. This rate was multiplied by 2.76 to get the 0-fold mutation rate, that is, the non-synonymous mutation rate, for PL. Note that the theta used for the SP population had to be constrained to theta PL × 0.74, to account for the difference in number of sites retained in each population after all filters for sequence quality. To estimate the DFE from the data collected for each gene group, we used a Poisson model including the population scaled mutation rate, theta, and compared the likelihood of the data along the parameter space. This model fits a DFE that is unbiased by mutations possibly too rare to be observed in the sample of genotypes used for computing the site frequency spectrum. Having determined the demographic parameters of the two populations and the DFE of each gene group, we proceeded to the third step of our analysis, which predicts the properties of genetic variation of each gene set and allows us to visualize the fit of the DFE and demography to the data. These properties follow from the DFE and demographic histories under the standard diffusion model. For this step, we calculated the distribution of selection coefficients for variants in each count of the SFS of each gene set. We first calculated the expected SFS for each selection coefficient under the demographic model using ∂a∂i functions. Then, we calculated the expected distribution of *s* using the python function gamma.cdf with the shape and scale parameter calculated for the joint estimate of the DFE of each gene set. Finally, we inferred the distribution of selection coefficients in each count of the SFS of each gene group by applying Bayes’ rule. The annotated code file we followed was provided as supplementary material in (Takou et al. 2021).

The confidence intervals for each group were based on values derived from 200 bootstraps obtained by resampling with replacement over genes. The statistical significance of differences between the groups in the DFE distribution was calculated by pairwise comparisons within each bin of the DFE (0-1, 1-10 and 1-inf). We used the union of the proportion obtained in all bootstraps to calculate what proportion of the mutations belong to the same group, multiplied by 2, to account for the fact that the test is one-sided (Keightley and Eyre-Walker 2007).

This method assumes that allelic effects are purely additive. If in fact variants had recessive effects on fitness, the method underestimates the magnitude of their effect. We note that this assumption should lead to a pattern opposite to the one reported in this manuscript, namely, variants in high-VA genes experienced weaker selection than variants in high-VNA genes. The results are therefore robust to the assumption of additive allelic effects on fitness.

### Quantification of amino-acid divergence

An alignment between *A. lyrata* and *A. thaliana,* which was produced in (He et al. 2021), was used to estimate the Ka and Ks values. The coding sequences of the orthologues genes were selected based on the genome annotation and aligned using MAFFT (Katoh and Standley 2013). The values were extracted by the function kaks of R package seqinr (Charif and Lobry 2007). The statistics were recalculated 1,000 times by bootstrap replicates of gene resampling with replacement.

### Genome-wide association study for identification of trans and cis regulatory effects on gene expression

We used the SNP calls of the 131 available mapped hybrid transcriptomes to identify variants associated with gene expression variation among families. We implemented a genome wide association study (GWAS) approach (Kang et al. 2010; Korte and Farlow 2013), with R packages downloaded from https://github.com/arthurkorte/GWAS. We filtered the SNP calls with vcftools to maintain only biallelic sites, with no missing information in any sample, minimum depth of 10 reads per sample, minimum SNP calling quality of 30, and minimum minor allele frequency of 0.05. The remaining 127,585 SNPs were used to estimate the kinship matrix with the function emma.kinship. The log-transformed counts of each of the 17,657 genes were successively used as the phenotypic input for each GWAS model (with separate models fit for each transcript). We retained the top hits for each GWAS using a Bonferoni cut off (*p* = 3.918956e-07) to control for multiple testing. Moreover, we grouped all individuals by their genotype at each detected SNP and computed the coefficient of variation (CV) of the tested gene’s expression between the two groups. CVs smaller than the 25^th^ percentile of the distribution were removed from the further analysis to exclude false positives.

## Acknowledgments

We would like to thank Kirsten Bell for help with the laboratory work, the Cologne Genome Center for the transcriptome sequences, and Emily Wheeler, Harwich, MA, USA, for editorial help. Furthermore, we would like to thank F. He, S. Sunyaev, O. Savolainen, T. Hämälä, V. Castric, A. Le Rouzic and M.G. Stetter for feedback, and F. Hospital for inspiring discussions ahead of this project. This research was funded by ERC Grant No. 648617 Adaptoscope.

## Data availability

The whole-genome and transcriptome sequences are available via ENA. The whole-genome sequences of the parental lines are under the number PRJEB33206, and the transcriptome sequences are under the number PRJEB49153. The gene counts per individual and the sql database compiling descriptors of genomic variation will be deposited in dryad upon acceptance of the manuscript. All the original scripts are in git repository https://github.com/mtakou/Quantitative-analysis-of-gene-expression.

