## Supplementary data for "Strength of purifying selection on the amino-acid sequence is associated with the amount of non-additive variance in gene expression"

**Takou et al. 2024**

### Supplementary Material

**Supplementary Table S1: Information about the full sibling families used in the analysis. Each full sibling family was generated based on the scheme presented in Figure 1.**

For each full sibling family, the Spiterstulen (SP) and the Plech (PL) sequence ID numbers, as described in Takou (2021) and annotated in ENA archive repository, are provided (PRJEB49153). For each family, the number of offspring used in the final analysis is shown.

| Family | SP parent | PL parent | Number of family members |
|---|---|---|---|
| Fam01 | 70539 | 10a | 11 |
| Fam02 | 70539 | 80936 | 18 |
| Fam03 | 70535 | 80936 | 13 |
| Fam04 | 70535 | 5a | 12 |
| Fam05 | 70537 | 6a | 9 |
| Fam06 | 70536 | 2a | 8 |
| Fam07 | 70544 | 6a | 20 |
| Fam08 | 70544 | 5a | 14 |
| Fam09 | 70545 | 2a | 6 |
| Fam10 | 70545 | 10a | 6 |
| Fam11 | 70536 | 11a | 0 |
| Fam12 | 70537 | 11a | 0 |
| Fam15 | 70539 | 2a | 4 |
| Fam16 | 70545 | 80936 | 12 |

Supplementary Table S2: **The p values for all pairwise comparisons within each DFE bin.**

The gene groups were specified based on their level of additive, dominance or genetic variance as well as on the sum of the residual and maternal variance.

| **$0 < N_{anc}s < 1$** | | | | |
|---|---|---|---|---|
| | **$V_A$** | **$V_{NA}$** | **$V_G$** | **$V_R$** |
| **Rand** | 0.03 | 0.3 | 0.01 | 0.01 |
| **$V_A$** | NA | 0.01 | 0.01 | 0.01 |
| **$V_D$** | | NA | 0.01 | 0.01 |
| **$V_G$** | | | NA | 0.01 |
| **$1 < N_{anc}s < 10$** | | | | |
| | **$V_A$** | **$V_{NA}$** | **$V_G$** | **$V_R$** |
| **Rand** | 0.23 | 0.19 | 0.01 | 0.01 |
| **Va** | NA | 0.52 | 0.01 | 0.01 |
| **Vd** | | NA | 0.01 | 0.01 |
| **$V_G$** | | | NA | 0.43 |
| **$10 < N_{anc}s < Inf$** | | | | |
| | **$V_A$** | **$V_{NA}$** | **$V_G$** | **$V_R$** |
| **Rand** | 0.01 | 0.39 | 0.01 | 0.01 |
| **$V_A$** | NA | 0.07 | 0.01 | 0.01 |
| **$V_D$** | | NA | 0.01 | 0.01 |
| **$V_G$** | | | NA | 0.01 |

Supplementary Table S3: **The importance, p -value, and significance of each predictor for $V_{NA}$ in the random forest analysis are provided.**

| motif | Importance | pvalue | sig |
|---|---|---|---|
| Exons | 53.021311278031 | 0.0163934426229508 | TRUE |
| Length | 26.9235437211514 | 0.0163934426229508 | TRUE |
| transcript.length | 14.2711690716638 | 0.0163934426229508 | TRUE |
| DOF5.3 | 4.72227694044062 | 0.0163934426229508 | TRUE |
| total | 4.34854187820101 | 0.0163934426229508 | TRUE |
| Density | 3.31778902806577 | 0.0163934426229508 | TRUE |
| AT3G24120 | 1.4476459603237 | 0.0163934426229508 | TRUE |
| AHL12 | 1.41265726405547 | 0.0163934426229508 | TRUE |
| AGL55 | 1.1773935987853 | 0.0163934426229508 | TRUE |
| ATHB.16 | 0.843748325988934 | 0.0163934426229508 | TRUE |
| GenesDEG | 0.824105253529609 | 0.0163934426229508 | TRUE |
| PL.pi | 0.724663289727124 | 0.0327868852459016 | TRUE |
| DOF2.4 | 0.593727808498788 | 0.0163934426229508 | TRUE |
| KAN1 | 0.533608889607024 | 0.0163934426229508 | TRUE |
| DOF5.6 | 0.529071143807268 | 0.0163934426229508 | TRUE |
| AGL42 | 0.469401111706553 | 0.0163934426229508 | TRUE |
| AHL20 | 0.292363578586661 | 0.0163934426229508 | TRUE |
| REF6 | 0.291366058737764 | 0.0163934426229508 | TRUE |
| PL.TajD | 0.27476288467216 | 0.147540983606557 | FALSE |
| dxy | 0.263116716456656 | 0.147540983606557 | FALSE |
| DOF1.8 | 0.248406050146087 | 0.0327868852459016 | TRUE |
| AT3G52440 | 0.246340663790506 | 0.0163934426229508 | TRUE |
| DOF5.7 | 0.245041317180332 | 0.0163934426229508 | TRUE |
| NAC080 | 0.227791514891445 | 0.0163934426229508 | TRUE |
| AHL25 | 0.213983496910369 | 0.0327868852459016 | TRUE |
| WRKY75 | 0.211819970215324 | 0.0327868852459016 | TRUE |
| NAC058 | 0.193050175006176 | 0.0163934426229508 | TRUE |
| SPL12 | 0.187553418624845 | 0.0327868852459016 | TRUE |
| MYB62 | 0.185649512806582 | 0.0163934426229508 | TRUE |
| ATHB.12 | 0.172691737394414 | 0.0327868852459016 | TRUE |
| OBP4 | 0.169116968133819 | 0.0163934426229508 | TRUE |
| ATHB.51 | 0.167174741144785 | 0.0819672131147541 | FALSE |
| NAC92 | 0.167045971113516 | 0.0163934426229508 | TRUE |
| ATHB13 | 0.155337886944448 | 0.0327868852459016 | TRUE |
| NAC083 | 0.154988042217836 | 0.0327868852459016 | TRUE |
| MYC4 | 0.141502982594132 | 0.0491803278688525 | TRUE |

| | | | |
|---|---|---|---|
| SPT | 0.138538828989848 | 0.0491803278688525 | TRUE |
| ABF3 | 0.135039573529432 | 0.0983606557377049 | FALSE |
| DAG2 | 0.130615750121603 | 0.0819672131147541 | FALSE |
| CCA1 | 0.127057404674366 | 0.0655737704918033 | FALSE |
| WRKY45 | 0.122930263481856 | 0.0327868852459016 | TRUE |
| GATA15 | 0.122253640485501 | 0.163934426229508 | FALSE |
| AT3G57600 | 0.11572014933352 | 0.0163934426229508 | TRUE |
| CDF2 | 0.109660570741975 | 0.163934426229508 | FALSE |
| ATHB18 | 0.101077721145036 | 0.0819672131147541 | FALSE |
| EDT1 | 0.100971348425886 | 0.0655737704918033 | FALSE |
| PIF4 | 0.0984163761307756 | 0.180327868852459 | FALSE |
| MYB119 | 0.0914462110160212 | 0.0819672131147541 | FALSE |
| WRKY62 | 0.091108708344051 | 0.0819672131147541 | FALSE |
| ATHB7 | 0.0904635359369356 | 0.0819672131147541 | FALSE |
| SPL3 | 0.0902790419554241 | 0.0983606557377049 | FALSE |
| FUS3 | 0.089846649818785 | 0.0655737704918033 | FALSE |
| ARR1 | 0.0890732288532928 | 0.0655737704918033 | FALSE |
| AT1G68670 | 0.0840135972932485 | 0.0655737704918033 | FALSE |
| BZR2 | 0.0756646983472359 | 0.0983606557377049 | FALSE |
| DOF2.5 | 0.0750556651234092 | 0.180327868852459 | FALSE |
| MYC3 | 0.0733342614860429 | 0.0655737704918033 | FALSE |
| bHLH69 | 0.0733328580842906 | 0.0327868852459016 | TRUE |
| bZIP68 | 0.0730785206357189 | 0.213114754098361 | FALSE |
| ARR14 | 0.0727144813276005 | 0.147540983606557 | FALSE |

| | | | |
|---|---|---|---|
| CDF3 | 0.0700649540322154 | 0.163934426229508 | FALSE |
| CMTA3 | 0.0683859077960443 | 0.327868852459016 | FALSE |
| AP1 | 0.068015704817839 | 0.163934426229508 | FALSE |
| ARR11 | 0.067371516460989 | 0.213114754098361 | FALSE |
| WRKY2 | 0.064981307366521 | 0.262295081967213 | FALSE |
| TCX2 | 0.0648416119735918 | 0.0655737704918033 | FALSE |
| HAT1 | 0.0619074296354447 | 0.180327868852459 | FALSE |
| STZ | 0.0613873766156475 | 0.229508196721311 | FALSE |
| CMTA2 | 0.060046450362535 | 0.229508196721311 | FALSE |
| AT2G03500 | 0.0598860639090205 | 0.262295081967213 | FALSE |
| SP.pi | 0.0575296187616815 | 0.262295081967213 | FALSE |
| AT1G76870 | 0.0564733259037383 | 0.229508196721311 | FALSE |
| ERF008 | 0.0564659547544929 | 0.245901639344262 | FALSE |
| At5g58900 | 0.054817869813478 | 0.0491803278688525 | TRUE |
| TCP5 | 0.0546685899893073 | 0.147540983606557 | FALSE |
| At5g08520 | 0.0540984310109035 | 0.131147540983607 | FALSE |
| ARF8 | 0.0527832225531968 | 0.278688524590164 | FALSE |
| AT4G36780 | 0.0506693441036999 | 0.0655737704918033 | FALSE |
| ERF1B | 0.0498966412496501 | 0.213114754098361 | FALSE |
| AT1G78700 | 0.0495880908915015 | 0.0819672131147541 | FALSE |
| ABI3 | 0.0491085008990543 | 0.39344262295082 | FALSE |
| BIM2 | 0.0483899108225 | 0.311475409836066 | FALSE |
| AT4G18890 | 0.0480543927623313 | 0.0491803278688525 | TRUE |

| | | | |
|---|---|---|---|
| ARF5 | 0.0479533872343772 | 0.213114754098361 | FALSE |
| At3g09600 | 0.0476747446592038 | 0.163934426229508 | FALSE |
| WRKY8 | 0.0473995818716582 | 0.180327868852459 | FALSE |
| HAT22 | 0.0465649381961525 | 0.229508196721311 | FALSE |
| AT1G47655 | 0.0460179425109655 | 0.295081967213115 | FALSE |
| WRKY15 | 0.0455382631719638 | 0.131147540983607 | FALSE |
| AT5G18450 | 0.0449205289095695 | 0.0983606557377049 | FALSE |
| TCP4 | 0.0448187614071357 | 0.213114754098361 | FALSE |
| MYB24 | 0.0446098218893268 | 0.213114754098361 | FALSE |
| SPL7 | 0.0441108722430062 | 0.327868852459016 | FALSE |
| ERF018 | 0.0438091203563471 | 0.344262295081967 | FALSE |
| MYB113 | 0.0408792021680905 | 0.344262295081967 | FALSE |
| AT2G40260 | 0.040158560791656 | 0.213114754098361 | FALSE |
| AT4G28140 | 0.0385913463396768 | 0.0655737704918033 | FALSE |
| ATHB53 | 0.0378383763732082 | 0.278688524590164 | FALSE |
| SOL1 | 0.0374644712851214 | 0.19672131147541 | FALSE |
| WRKY23 | 0.0371154802886182 | 0.344262295081967 | FALSE |
| BHLH34 | 0.0369597260439536 | 0.278688524590164 | FALSE |
| ATHB33 | 0.0355335217080269 | 0.360655737704918 | FALSE |
| RAP26 | 0.0354782003878207 | 0.0655737704918033 | FALSE |
| WRKY18 | 0.034114957248133 | 0.245901639344262 | FALSE |
| RAP211 | 0.0328915207992887 | 0.229508196721311 | FALSE |
| UNE10 | 0.0326426526510559 | 0.278688524590164 | FALSE |

| | | | |
|---|---|---|---|
| ATHB24 | 0.0325542784923487 | 0.229508196721311 | FALSE |
| TGA10 | 0.0324730451572742 | 0.147540983606557 | FALSE |
| DRE1C | 0.03242297974909 | 0.327868852459016 | FALSE |
| WRKY38 | 0.0308316021313298 | 0.295081967213115 | FALSE |
| SPL9 | 0.0306701476362694 | 0.295081967213115 | FALSE |
| AT3G10580 | 0.030466091513631 | 0.131147540983607 | FALSE |
| AIB | 0.0275612792784537 | 0.0491803278688525 | TRUE |
| ANL2 | 0.0273847955192974 | 0.278688524590164 | FALSE |
| WRKY63 | 0.0269551471338648 | 0.278688524590164 | FALSE |
| ARR10 | 0.0268330427594155 | 0.360655737704918 | FALSE |
| DREB1E | 0.026341189953408 | 0.442622950819672 | FALSE |
| TINY | 0.0254753688121996 | 0.0655737704918033 | FALSE |
| BEE2 | 0.0247983106781785 | 0.163934426229508 | FALSE |
| TGA5 | 0.0243538687047103 | 0.327868852459016 | FALSE |
| BHLH78 | 0.0237669039600667 | 0.0491803278688525 | TRUE |
| WRKY48 | 0.0230038266601833 | 0.377049180327869 | FALSE |
| WRKY25 | 0.0225841608370838 | 0.295081967213115 | FALSE |
| WRKY43 | 0.0216152325194314 | 0.245901639344262 | FALSE |
| RAV1.var.2. | 0.02138835380085 | 0.262295081967213 | FALSE |
| MYB3R4 | 0.0212535621038254 | 0.147540983606557 | FALSE |
| PIF1 | 0.0202004754603495 | 0.262295081967213 | FALSE |
| MYB98 | 0.0195069351878718 | 0.262295081967213 | FALSE |
| TCP16 | 0.0189785211213533 | 0.360655737704918 | FALSE |

| | | | |
|---|---|---|---|
| dof4.5 | 0.0187407230062742 | 0.311475409836066 | FALSE |
| BPC6 | 0.0179992290348061 | 0.442622950819672 | FALSE |
| WRKY12 | 0.0168980448539437 | 0.262295081967213 | FALSE |
| WRKY59 | 0.0168938140671882 | 0.475409836065574 | FALSE |
| BHLH3 | 0.016885233919142 | 0.180327868852459 | FALSE |
| PIF5 | 0.0168781397317865 | 0.360655737704918 | FALSE |
| At5g47390 | 0.0166372802174385 | 0.213114754098361 | FALSE |
| CRF2 | 0.0157497959600738 | 0.39344262295082 | FALSE |
| CBF2 | 0.0154942808553289 | 0.311475409836066 | FALSE |
| WRKY22 | 0.0147381874918868 | 0.245901639344262 | FALSE |
| GenesSweep | 0.0146483122021537 | 0.360655737704918 | FALSE |
| TGA3 | 0.0145243001938187 | 0.229508196721311 | FALSE |
| KAN4 | 0.013969327531154 | 0.491803278688525 | FALSE |
| WRKY28 | 0.0131111163536076 | 0.213114754098361 | FALSE |
| ATHB25 | 0.0129306575549607 | 0.426229508196721 | FALSE |
| ATHB20 | 0.0121825516315361 | 0.459016393442623 | FALSE |
| MYB3R1 | 0.0111920868406973 | 0.360655737704918 | FALSE |
| SPL5 | 0.0109551575434286 | 0.344262295081967 | FALSE |
| TSO1 | 0.0108631190948756 | 0.327868852459016 | FALSE |
| AGL16 | 0.0107391934768 | 0.344262295081967 | FALSE |
| SPL14 | 0.0104582082159664 | 0.344262295081967 | FALSE |
| bHLH130 | 0.00992362027088798 | 0.409836065573771 | FALSE |
| HDG1.100_a | 0.00979187970582301 | 0.459016393442623 | FALSE |

| | | | |
|---|---|---|---|
| WRKY21 | 0.00972008817731831 | 0.508196721311475 | FALSE |
| BHLH13 | 0.00967558485859062 | 0.213114754098361 | FALSE |
| PI | 0.00868044153440669 | 0.377049180327869 | FALSE |
| WRKY71 | 0.00867751076125095 | 0.278688524590164 | FALSE |
| ZAP1 | 0.00755746005174986 | 0.442622950819672 | FALSE |
| AT3G16280 | 0.00754992819393434 | 0.278688524590164 | FALSE |
| ATHB.6 | 0.00744616173131946 | 0.39344262295082 | FALSE |
| LCL1 | 0.00739113947958582 | 0.426229508196721 | FALSE |
| AT1G22810 | 0.00710258892380681 | 0.426229508196721 | FALSE |
| Adof1 | 0.00688941528765647 | 0.131147540983607 | FALSE |
| dof4.2 | 0.0067872978664105 | 0.311475409836066 | FALSE |
| MGP | 0.00643640778404648 | 0.0819672131147541 | FALSE |
| AP3 | 0.00640709248212543 | 0.508196721311475 | FALSE |
| AT5G29000 | 0.00633255068167355 | 0.163934426229508 | FALSE |
| AGL13 | 0.00608923808772694 | 0.147540983606557 | FALSE |
| ESE1 | 0.00575305293421963 | 0.459016393442623 | FALSE |
| ABI5 | 0.0053536848755132 | 0.557377049180328 | FALSE |
| PIF7 | 0.00500769745839321 | 0.0163934426229508 | TRUE |
| WRKY26 | 0.00499237670849806 | 0.39344262295082 | FALSE |
| IDD2 | 0.00487449410143276 | 0.459016393442623 | FALSE |
| BZIP60 | 0.00451888463849577 | 0.377049180327869 | FALSE |
| At5g52660 | 0.00450688841060273 | 0.508196721311475 | FALSE |
| ERF4 | 0.00449884072091597 | 0.491803278688525 | FALSE |

| | | | |
|---|---|---|---|
| ERF38 | 0.0041037580780665 | 0.262295081967213 | FALSE |
| NAC043 | 0.00392522098025757 | 0.573770491803279 | FALSE |
| AT4G16750 | 0.00378522624635454 | 0.295081967213115 | FALSE |
| TGA6 | 0.00374729063151495 | 0.442622950819672 | FALSE |
| bZIP52 | 0.00338590814814634 | 0.442622950819672 | FALSE |
| At5g05790 | 0.00300668225351897 | 0.491803278688525 | FALSE |
| ERF5 | 0.00277841628886928 | 0.459016393442623 | FALSE |
| COG1 | 0.00247057089212932 | 0.459016393442623 | FALSE |
| DEAR3 | 0.00244331471951947 | 0.426229508196721 | FALSE |
| bHLH77 | 0.00230175251110822 | 0.426229508196721 | FALSE |
| TGA4 | 0.00226655305855452 | 0.442622950819672 | FALSE |
| WRKY24 | 0.00223557285929196 | 0.377049180327869 | FALSE |
| MYC2 | 0.00186883696624919 | 0.442622950819672 | FALSE |
| WRKY14 | 0.00184147029757332 | 0.311475409836066 | FALSE |
| GT2 | 0.00178966035986669 | 0.426229508196721 | FALSE |
| AGL3 | 0.00156160052430158 | 0.409836065573771 | FALSE |
| AT2G33710 | 0.00135291137856128 | 0.409836065573771 | FALSE |
| CBF1 | 0.00122940019368704 | 0.475409836065574 | FALSE |
| AGL27 | 0.00107244017060185 | 0.426229508196721 | FALSE |
| GT3a | 0.000777769200248468 | 0.213114754098361 | FALSE |
| MYB57 | 0.000762093859790394 | 0.540983606557377 | FALSE |
| AGL15 | 0.00050314327792178 | 0.295081967213115 | FALSE |
| AG | 0.000499334011531237 | 0.344262295081967 | FALSE |

| | | | | |
|---|---|---|---|---|
| Sep-01 | 0.000464802928128693 | 0.508196721311475 | | FALSE |
| JKD | 0.000320531286413803 | 0.360655737704918 | | FALSE |
| ERF9 | 0.000307093292000228 | 0.163934426229508 | | FALSE |
| AT5G66940 | 0.000283744531063409 | 0.229508196721311 | | FALSE |
| RAP212 | 0.000180969520996906 | 0.442622950819672 | | FALSE |
| IDD5 | 0.000159776022482217 | 0.409836065573771 | | FALSE |
| SVP | 0.000119108560767098 | 0.491803278688525 | | FALSE |
| ATHB23 | 9,16E+09 | 0.491803278688525 | | FALSE |
| LEP | 4,47E+09 | 0.426229508196721 | | FALSE |
| AGL25 | 1,72E+09 | 0.377049180327869 | | FALSE |
| DREB19 | 0 | 0.688524590163934 | | FALSE |
| bHLH74 | 0 | 1 | | FALSE |
| AT1G12630 | 0 | 1 | | FALSE |
| TCP1 | 0 | 1 | | FALSE |
| AGL1 | 0 | 0.590163934426229 | | FALSE |
| NUC | 0 | 1 | | FALSE |
| AT3G25990 | 0 | 1 | | FALSE |
| MYB15 | 0 | 1 | | FALSE |
| At5g08330 | -1,12E+09 | 0.622950819672131 | | FALSE |
| AGL6 | -4,05E+09 | 0.60655737704918 | | FALSE |
| AT3G45610 | -0.000140895826769092 | 0.475409836065574 | | FALSE |
| ABR1 | -0.000199145284101136 | 0.754098360655738 | | FALSE |

| | | | |
|---|---|---|---|
| MYB56 | -0.000205957694237913 | 0.377049180327869 | FALSE |
| SPL15 | -0.000295263389901723 | 0.60655737704918 | FALSE |
| At1g72010 | -0.000312975966817282 | 0.60655737704918 | FALSE |
| HBI1 | -0.000388270316845363 | 0.557377049180328 | FALSE |
| TCP14 | -0.000444326105097695 | 0.786885245901639 | FALSE |
| WRKY6 | -0.000608565462110181 | 0.672131147540984 | FALSE |
| HY5 | -0.000622863767942647 | 0.508196721311475 | FALSE |
| bHLH18 | -0.000868764287886619 | 0.688524590163934 | FALSE |
| AT1G13300 | -0.000888456942651474 | 0.622950819672131 | FALSE |
| BZR1 | -0.000949101588682136 | 0.868852459016393 | FALSE |
| TRP2 | -0.00100797438886625 | 0.524590163934426 | FALSE |
| WRKY27 | -0.00111839851680507 | 0.491803278688525 | FALSE |
| bZIP50 | -0.00115874971916208 | 0.639344262295082 | FALSE |
| FHY3 | -0.00135047392834604 | 0.557377049180328 | FALSE |
| AT4G18450 | -0.00135826876296831 | 0.459016393442623 | FALSE |

| | | | |
|---|---|---|---|
| WRKY11 | -0.00140207560116523 | 0.557377049180328 | FALSE |
| ERF098 | -0.00169767840137155 | 0.60655737704918 | FALSE |
| AT3G12730 | -0.00179094068242337 | 0.377049180327869 | FALSE |
| DREB26 | -0.00179861235279404 | 0.639344262295082 | FALSE |
| ESE3 | -0.00245938614370692 | 0.508196721311475 | FALSE |
| KAN2 | -0.00251252661079676 | 0.60655737704918 | FALSE |
| RAV1 | -0.00264658128704366 | 0.540983606557377 | FALSE |
| ERF2 | -0.00286508844134382 | 0.672131147540984 | FALSE |
| ERF104 | -0.00308792006911622 | 0.573770491803279 | FALSE |
| TRP1 | -0.00325455085248791 | 0.655737704918033 | FALSE |
| AT1G69570 | -0.00329361625980444 | 0.639344262295082 | FALSE |
| ERF15 | -0.00332447409556176 | 0.557377049180328 | FALSE |
| bZIP53 | -0.00357240849096749 | 0.639344262295082 | FALSE |
| MYB27 | -0.00357424172738944 | 0.590163934426229 | FALSE |
| TGA9 | -0.0036188381088787 | 0.672131147540984 | FALSE |

| | | | |
|---|---|---|---|
| TCP19 | -0.00367026365354358 | 0.590163934426229 | FALSE |
| AT1G71450 | -0.00369780948687624 | 0.770491803278688 | FALSE |
| MYB46 | -0.00386859327841282 | 0.573770491803279 | FALSE |
| ATHB40 | -0.00401481448753917 | 0.80327868852459 | FALSE |
| IDD4 | -0.00430539260831877 | 0.704918032786885 | FALSE |
| AT4G12670 | -0.00436420305414258 | 0.80327868852459 | FALSE |
| WRKY42 | -0.00441030164535087 | 0.868852459016393 | FALSE |
| CDC5 | -0.00452105865611751 | 0.557377049180328 | FALSE |
| ERF10 | -0.00453570725207858 | 0.704918032786885 | FALSE |
| WRKY3 | -0.00471505577404064 | 0.590163934426229 | FALSE |
| E2FA | -0.00512694238662848 | 0.573770491803279 | FALSE |
| AT5G56840 | -0.00568082032413415 | 0.639344262295082 | FALSE |
| MYB59 | -0.00590363933119687 | 0.540983606557377 | FALSE |
| ERF13 | -0.00616314987157851 | 0.590163934426229 | FALSE |
| BPC5 | -0.00653256181489651 | 0.540983606557377 | FALSE |

| | | | | |
|---|---|---|---|---|
| WRKY46 | -0.00703890910309764 | 0.60655737704918 | | FALSE |
| PIF3 | -0.00741024871015636 | 0.60655737704918 | | FALSE |
| AT1G76880 | -0.00750625751009311 | 0.622950819672131 | | FALSE |
| WRKY17 | -0.00757624900902938 | 0.80327868852459 | | FALSE |
| MYB118 | -0.00779901269471062 | 0.885245901639344 | | FALSE |
| WRKY65 | -0.00783260165782744 | 0.655737704918033 | | FALSE |
| ATHB.5 | -0.00788572162548553 | 0.704918032786885 | | FALSE |
| OBP1 | -0.00799800357322313 | 0.508196721311475 | | FALSE |
| AT1G28160 | -0.00835553532347185 | 0.737704918032787 | | FALSE |
| WRKY33 | -0.00856741359175106 | 0.639344262295082 | | FALSE |
| AT1G36060 | -0.0085842716493759 | 0.737704918032787 | | FALSE |
| DEAR5 | -0.00898816599790475 | 0.721311475409836 | | FALSE |
| MYB3R5 | -0.00966762738962615 | 0.672131147540984 | | FALSE |
| AT4G32800 | -0.00968856736376286 | 0.704918032786885 | | FALSE |
| BIM3 | -0.00985249851373859 | 0.573770491803279 | | FALSE |

| | | | | |
|---|---|---|---|---|
| AREB3 | -0.010588669910001 | 0.606557377049 18 | | FALSE |
| WRKY70 | -0.010634898801062 | 0.770491803278688 | | FALSE |
| AT3G46070 | -0.0108101183321198 | 0.721311475409836 | | FALSE |
| ERF3 | -0.0114716721938325 | 0.590163934426229 | | FALSE |
| WRKY29 | -0.0117298034318914 | 0.655737704918033 | | FALSE |
| AT2G44940 | -0.0117307463431737 | 0.655737704918033 | | FALSE |
| TGA7 | -0.0124048571269861 | 0.590163934426229 | | FALSE |
| GBF6 | -0.0126642457044153 | 0.737704918032787 | | FALSE |
| ATHB21 | -0.0127770415209166 | 0.721311475409836 | | FALSE |
| PUCHI | -0.0128481049166986 | 0.737704918032787 | | FALSE |
| HAT2 | -0.0130321030132838 | 0.655737704918033 | | FALSE |
| ERF7 | -0.0130559806118158 | 0.557377049180328 | | FALSE |
| At1g74840 | -0.0141792693474478 | 0.639344262295082 | | FALSE |
| bZIP48 | -0.0141938215161427 | 0.688524590163934 | | FALSE |
| SOC1 | -0.0145032596822145 | 0.737704918032787 | | FALSE |
| GATA14 | -0.0146120589177539 | 0.803278688524 59 | | FALSE |

| | | | |
|---|---|---|---|
| LHY1 | -0.0147965328299624 | 0.655737704918033 | FALSE |
| AT1G25550 | -0.0155230687893276 | 0.540983606557377 | FALSE |
| ARF2 | -0.0157237125935875 | 0.639344262295082 | FALSE |
| bZIP44 | -0.0158768931551765 | 0.868852459016393 | FALSE |
| At3g11280 | -0.0162590536573994 | 0.622950819672131 | FALSE |
| AIL7 | -0.0163840790595437 | 0.655737704918033 | FALSE |
| bZIP3 | -0.016670223228793 | 0.868852459016393 | FALSE |
| ERF8 | -0.0171720837318292 | 0.60655737704918 | FALSE |
| ATHB15 | -0.0176462876486955 | 0.885245901639344 | FALSE |
| ERF105 | -0.0176542597890031 | 0.737704918032787 | FALSE |
| RAP210 | -0.0178052430847505 | 0.737704918032787 | FALSE |
| bZIP43 | -0.017838822809641 | 0.819672131147541 | FALSE |
| AT1G19210 | -0.0179367994070447 | 0.655737704918033 | FALSE |
| ERF069 | -0.0179540952202288 | 0.672131147540984 | FALSE |
| At1g49010 | -0.0181485303180951 | 0.655737704918033 | FALSE |
| TCP17 | -0.0183738094896929 | 0.754098360655738 | FALSE |

| | | | |
|---|---|---|---|
| FAR1 | -0.0193057077819644 | 0.672131147540984 | FALSE |
| AT2G20110 | -0.0195431377754046 | 0.688524590163934 | FALSE |
| AT5G45580 | -0.0197052769182499 | 0.622950819672131 | FALSE |
| TBP3 | -0.0197301316439427 | 0.60655737704918 | FALSE |
| AT1G75490 | -0.0198306228405159 | 0.819672131147541 | FALSE |
| OBP3 | -0.0198435624483268 | 0.721311475409836 | FALSE |
| bHLH80 | -0.0201437458143006 | 0.557377049180328 | FALSE |
| MYB70 | -0.0205477385133589 | 0.721311475409836 | FALSE |
| WRKY7 | -0.0206807472519658 | 0.918032786885246 | FALSE |
| ARR18 | -0.0207891437356831 | 0.672131147540984 | FALSE |
| MYB3 | -0.0208208645329842 | 0.672131147540984 | FALSE |
| WRKY57 | -0.0212100058405499 | 0.754098360655738 | FALSE |
| DYT1 | -0.0212283059350595 | 0.901639344262295 | FALSE |
| WRKY31 | -0.0217408827646114 | 0.934426229508197 | FALSE |
| TCP2 | -0.0218149826845052 | 0.950819672131147 | FALSE |

| | | | |
|---|---|---|---|
| IDD7 | -0.0222907069114607 | 0.934426229508197 | FALSE |
| AT1G44830 | -0.022677197679523 | 0.786885245901639 | FALSE |
| GBF3 | -0.0230975482805446 | 0.852459016393443 | FALSE |
| MYB4 | -0.023465267500035 | 0.655737704918033 | FALSE |
| NTL9 | -0.0243769423436388 | 0.704918032786885 | FALSE |
| WRKY47 | -0.024513431327516 | 0.934426229508197 | FALSE |
| MYB73 | -0.0246611208310519 | 0.655737704918033 | FALSE |
| MYB101 | -0.0246807985334035 | 0.786885245901639 | FALSE |
| AT5G61620 | -0.0249619530380063 | 0.868852459016393 | FALSE |
| AT2G01060 | -0.0251426988107101 | 0.770491803278688 | FALSE |
| AT2G28810 | -0.0260237708771022 | 0.885245901639344 | FALSE |
| MYB111 | -0.0265173965031409 | 0.639344262295082 | FALSE |
| PTF1 | -0.0291083542365339 | 0.967213114754098 | FALSE |
| At1g19000 | -0.029199538055655 | 0.852459016393443 | FALSE |
| AT3G60490 | -0.029369754061825 | 0.885245901639344 | FALSE |
| ERF096 | -0.029455013323573 | 0.721311475409836 | FALSE |
| AT1G01250 | -0.0306760208574611 | 0.934426229508197 | FALSE |

| | | | |
|---|---|---|---|
| GATA10 | -0.0310808721938404 | 0.819672131147541 | FALSE |
| MYB105 | -0.0318893321470221 | 0.754098360655738 | FALSE |
| ABF2 | -0.0322382638827867 | 0.721311475409836 | FALSE |
| TCP20 | -0.0323520068129837 | 0.836065573770492 | FALSE |
| EPR1 | -0.0343992873357001 | 0.721311475409836 | FALSE |
| Sep-03 | -0.0344459019963886 | 0.754098360655738 | FALSE |
| LEC2 | -0.0344712758647899 | 0.688524590163934 | FALSE |
| RVE1 | -0.034549540526883 | 0.80327868852459 | FALSE |
| bZIP42 | -0.0358046828288628 | 0.950819672131147 | FALSE |
| MYB55 | -0.0362166705333427 | 0.737704918032787 | FALSE |
| DREB1A | -0.0371330234656047 | 0.868852459016393 | FALSE |
| TCP7 | -0.0376635464013562 | 0.901639344262295 | FALSE |
| SPL8 | -0.0384061703458114 | 0.754098360655738 | FALSE |
| GATA20 | -0.0384257395788297 | 0.721311475409836 | FALSE |
| GATA9 | -0.0385995899520592 | 0.688524590163934 | FALSE |
| bZIP16 | -0.040153196174927 | 0.852459016393443 | FALSE |

| | | | |
|---|---|---|---|
| DREB2C | -0.0404272127319666 | 0.770491803278688 | FALSE |
| AT4G37180 | -0.0411001253433473 | 0.737704918032787 | FALSE |
| GATA12 | -0.0419210531606982 | 0.672131147540984 | FALSE |
| RAP2.3 | -0.0424028302487021 | 0.967213114754098 | FALSE |
| GATA19 | -0.0429260818377503 | 0.80327868852459 | FALSE |
| CEJ1 | -0.0438391257677284 | 0.901639344262295 | FALSE |
| AGL63 | -0.044068530862193 | 0.934426229508197 | FALSE |
| WRKY60 | -0.0465685286454465 | 0.918032786885246 | FALSE |
| AT5G47660 | -0.0471674494872831 | 0.885245901639344 | FALSE |
| bZIP28 | -0.0478770293731296 | 0.934426229508197 | FALSE |
| MYB65 | -0.04874220618484 | 0.868852459016393 | FALSE |
| At4g01280 | -0.0487899993827113 | 0.967213114754098 | FALSE |
| WRKY55 | -0.050262710869214 | 0.918032786885246 | FALSE |
| BHLH104 | -0.0507021261910605 | 0.934426229508197 | FALSE |
| ATHB4 | -0.0511313497852257 | 0.688524590163934 | FALSE |
| SPL13 | -0.0520554314699608 | 0.868852459016393 | FALSE |

| | | | |
|---|---|---|---|
| SPL1 | -0.0520723289235723 | 0.901639344262295 | FALSE |
| ARF3 | -0.0529141810547167 | 0.704918032786885 | FALSE |
| ERF043 | -0.0537224227666152 | 0.885245901639344 | FALSE |
| RAP2.6 | -0.054907476956152 | 0.983606557377049 | FALSE |
| SPL4 | -0.0557300863748839 | 0.819672131147541 | FALSE |
| SGR5 | -0.0558510071092026 | 0.754098360655738 | FALSE |
| ERF094 | -0.0565965208678114 | 0.754098360655738 | FALSE |
| CAMTA1 | -0.0566653556965319 | 0.918032786885246 | FALSE |
| BPC1 | -0.0575981401071717 | 0.770491803278688 | FALSE |
| AT5G02460 | -0.0592020729078986 | 0.885245901639344 | FALSE |
| T11I18.17 | -0.0594520249870048 | 0.80327868852459 | FALSE |
| ERF109 | -0.059699611245122 | 0.967213114754098 | FALSE |
| RAP2.10 | -0.0601017974724582 | 1 | FALSE |
| GT.1 | -0.0611772513931021 | 0.737704918032787 | FALSE |
| AT3G10113 | -0.0618075755598018 | 0.934426229508197 | FALSE |
| NAC025 | -0.0631683137366167 | 0.770491803278688 | FALSE |

| | | | |
|---|---|---|---|
| ERF11 | -0.0641040002739932 | 0.918032786885246 | FALSE |
| AT1G72740 | -0.0654352097910387 | 0.852459016393443 | FALSE |
| TRB2 | -0.0657431933784094 | 0.80327868852459 | FALSE |
| MYB77 | -0.0666102790923489 | 0.950819672131147 | FALSE |
| HAT5 | -0.0667065343380642 | 0.967213114754098 | FALSE |
| TGA2 | -0.0679916569118177 | 0.836065573770492 | FALSE |
| F3A4.140 | -0.0707792985302501 | 0.918032786885246 | FALSE |
| WRKY20 | -0.0712800763689822 | 1 | FALSE |
| MYB33 | -0.0727190890508577 | 0.885245901639344 | FALSE |
| ERF112 | -0.0741421776002923 | 0.950819672131147 | FALSE |
| WRKY30 | -0.0742315910518428 | 0.934426229508197 | FALSE |
| CRF4 | -0.0747862300469551 | 1 | FALSE |
| MYB52 | -0.0761972623222427 | 0.819672131147541 | FALSE |
| WRKY40 | -0.0766664299234801 | 0.836065573770492 | FALSE |
| TCP3 | -0.0771008311637724 | 1 | FALSE |

| | | | |
|---|---|---|---|
| ATHB34 | -0.077873653291783 | 0.885245901639344 | FALSE |
| RAP21 | -0.0810695790281678 | 0.967213114754098 | FALSE |
| TCP24 | -0.0812087349801031 | 0.950819672131147 | FALSE |
| AT1G49560 | -0.0852499433044093 | 0.885245901639344 | FALSE |
| GATA11 | -0.085605807063064 | 0.950819672131147 | FALSE |
| TCP23 | -0.0856526996123633 | 0.967213114754098 | FALSE |
| SPL11 | -0.0857321917356486 | 0.967213114754098 | FALSE |
| AT1G77200 | -0.0858446953094304 | 0.983606557377049 | FALSE |
| BIM1 | -0.0875266296092996 | 0.934426229508197 | FALSE |
| CBF4 | -0.0909839414866514 | 0.983606557377049 | FALSE |
| WRKY50 | -0.091297909574517 | 1 | FALSE |
| SMZ | -0.091614461999637 | 0.852459016393443 | FALSE |
| TGA1 | -0.092191966526813 | 0.950819672131147 | FALSE |
| ERF039 | -0.0939803816907029 | 0.918032786885246 | FALSE |
| AT5G67000 | -0.0944190136058652 | 0.950819672131147 | FALSE |
| TCP15 | -0.0969808696365993 | 0.983606557377049 | FALSE |
| GATA8 | -0.114775971820281 | 0.819672131147541 | FALSE |

| | | | |
|---|---|---|---|
| UIF1 | -0.117320555182519 | 0.934426229508197 | FALSE |
| ERF6 | -0.121692852091675 | 0.983606557377049 | FALSE |
| AT3G04030 | -0.12216483907351 | 0.950819672131147 | FALSE |
| NAC055 | -0.12313265357609 | 0.934426229508197 | FALSE |
| MYB81 | -0.127905756642888 | 0.983606557377049 | FALSE |
| ARR2 | -0.131899653202723 | 1 | FALSE |
| ARF1 | -0.133606609621449 | 0.934426229508197 | FALSE |
| GATA6 | -0.139341326886418 | 1 | FALSE |
| Fst | -0.241558739160199 | 0.754098360655738 | FALSE |
| SP.TajD | -0.577513629749457 | 0.967213114754098 | FALSE |

**Supplementary Table S4: Correlation of dominance and additive variance with genome architecture traits and summary statistics of population genetics.** For each correlation between the fraction of variance and the genomic feature, the p value, Spearman's ρ as well as the number of cluster genes that were grouped are given. When the number of clusters is 1, the expression level of each gene was treated as independent observation.

| **Variance** | **Feature** | **Number of clusters** | ***P* value** | **Spearman's ρ** |
|---|---|---|---|---|
| Non Additive | Transcript Length | 1 | 1.22e-266 | 0.2588 |
| | Median Gene Expression | 1 | 9.54e-16 | 0.093 |
| | Number of Exons | 1 | <2.2e-16 | 0.3852 |
| | Gene Number | 200 | 1.253e-13 | 0.49 |
| | Transcript Length | 200 | 0.0042 | 0.201 |
| | Number of Exons | 200 | 7.752e-11 | 0.439 |
| Additive | Transcript Length | 1 | <2.2e-16 | -0.177 |
| | $F_{ST}$ | 1 | 0.0152 | -0.023 |
| | Dxy | 1 | <2.2e-16 | 0.1152 |
| | π of SP | 1 | <2.2e-16 | 0.0916 |
| | π of PL | 1 | <2.2e-16 | 0.115 |
| | Gene Number | 200 | 1.101e-05 | -0.277 |
| | $F_{ST}$ | 200 | 0.0041 | -0.201 |
| | Dxy | 200 | 2.59e-06 | 0.32 |
| | π of SP | 200 | 8.643e-07 | 0.34 |
| | π of PL | 200 | 1.627e-07 | 0.33 |

Supplementary Table S5: Gene ontology enrichment analysis among genes with high non-additive variance $V_{NA}$. The GOs ID, the term and the p value of the significant GOs are given. We used permutations to determine a conservative p value threshold (p<0.038) below which GO enrichments were not detected in a randomly ranked set of genes.

| GO.ID | Term | *P* value |
|---|---|---|
| GO:0031048 | chromatin silencing by small RNA | 4.5e-12 |
| GO:0000911 | cytokinesis by cell plate formation | 2.0e-11 |
| GO:0006306 | DNA methylation | 4.3e-11 |
| GO:0000956 | nuclear-transcribed mRNA catabolic proce... | 9.2e-11 |
| GO:0006346 | methylation-dependent chromatin silencin... | 6.3e-10 |
| GO:0008150 | biological_process | 2.1e-09 |
| GO:0045010 | actin nucleation | 3.5e-09 |
| GO:0010090 | trichome morphogenesis | 5.6e-09 |
| GO:0051567 | histone H3-K9 methylation | 1.6e-08 |
| GO:0007062 | sister chromatid cohesion | 1.8e-08 |
| GO:0009630 | gravitropism | 3.1e-08 |
| GO:0035196 | production of miRNAs involved in gene si... | 1.3e-07 |
| GO:0007131 | reciprocal meiotic recombination | 7.9e-07 |
| GO:0051225 | spindle assembly | 8.8e-07 |
| GO:0006275 | regulation of DNA replication | 1.1e-06 |
| GO:0010267 | production of ta-siRNAs involved in RNA ... | 1.2e-06 |
| GO:0009616 | virus induced gene silencing | 2.0e-06 |
| GO:0016572 | histone phosphorylation | 3.4e-06 |
| GO:0016558 | protein import into | 3.7e-06 |

| | | |
|---|---|---|
| | peroxisome matrix | |
| GO:0045132 | meiotic chromosome segregation | 4.8e-06 |
| GO:0009909 | regulation of flower development | 5.4e-06 |
| GO:0000278 | mitotic cell cycle | 5.5e-06 |
| GO:0007267 | cell-cell signaling | 8.5e-06 |
| GO:0000724 | double-strand break repair via homologous... | 9.0e-06 |
| GO:0007129 | synapsis | 1.1e-05 |
| GO:0015996 | chlorophyll catabolic process | 1.9e-05 |
| GO:0007155 | cell adhesion | 2.1e-05 |
| GO:0000226 | microtubule cytoskeleton organization | 2.6e-05 |
| GO:0006635 | fatty acid beta-oxidation | 2.7e-05 |
| GO:0006270 | DNA replication initiation | 4.0e-05 |
| GO:0032204 | regulation of telomere maintenance | 6.2e-05 |
| GO:0046855 | inositol phosphate dephosphorylation | 6.4e-05 |
| GO:0043247 | telomere maintenance in response to DNA ... | 7.0e-05 |
| GO:0010228 | vegetative to reproductive phase transit... | 7.8e-05 |
| GO:0010389 | regulation of G2/M transition of mitotic... | 8.5e-05 |
| GO:0042138 | meiotic DNA double-strand break formatio... | 8.9e-05 |
| GO:0032957 | inositol trisphosphate metabolic process | 9.1e-05 |

| | | |
|---|---|---|
| GO:0010332 | response to gamma radiation | 9.1e-05 |
| GO:0008283 | cell proliferation | 0.00018 |
| GO:0009855 | determination of bilateral symmetry | 0.00025 |
| GO:0043687 | post-translational protein modification | 0.00026 |
| GO:0016926 | protein desumoylation | 0.00044 |
| GO:0006342 | chromatin silencing | 0.00046 |
| GO:0008380 | RNA splicing | 0.00050 |
| GO:0006623 | protein targeting to vacuole | 0.00057 |
| GO:0010264 | myo-inositol hexakisphosphate biosynthet... | 0.00062 |
| GO:0006312 | mitotic recombination | 0.00065 |
| GO:0006486 | protein glycosylation | 0.00067 |
| GO:0016444 | somatic cell DNA recombination | 0.00089 |
| GO:0010638 | positive regulation of organelle organiz... | 0.00099 |
| GO:0051276 | chromosome organization | 0.00113 |
| GO:0009887 | animal organ morphogenesis | 0.00120 |
| GO:0016246 | RNA interference | 0.00126 |
| GO:0006406 | mRNA export from nucleus | 0.00135 |
| GO:0043414 | macromolecule methylation | 0.00146 |
| GO:0008284 | positive regulation of cell proliferatio... | 0.00156 |
| GO:0006606 | protein import into nucleus | 0.00158 |
| GO:0010014 | meristem initiation | 0.00171 |
| GO:0000280 | nuclear division | 0.00201 |
| GO:0032875 | regulation of DNA endoreduplication | 0.00202 |

| | | |
|---|---|---|
| GO:0006665 | sphingolipid metabolic process | 0.00214 |
| GO:0010050 | vegetative phase change | 0.00229 |
| GO:0043543 | protein acylation | 0.00245 |
| GO:0050665 | hydrogen peroxide biosynthetic process | 0.00270 |
| GO:0016192 | vesicle-mediated transport | 0.00288 |
| GO:0006487 | protein N-linked glycosylation | 0.00296 |
| GO:0009640 | photomorphogenesis | 0.00338 |
| GO:0048573 | photoperiodism, flowering | 0.00357 |
| GO:0030029 | actin filament-based process | 0.00362 |
| GO:0006397 | mRNA processing | 0.00364 |
| GO:0007346 | regulation of mitotic cell cycle | 0.00383 |
| GO:0010564 | regulation of cell cycle process | 0.00400 |
| GO:0031047 | gene silencing by RNA | 0.00400 |
| GO:0048449 | floral organ formation | 0.00419 |
| GO:0016570 | histone modification | 0.00441 |
| GO:0009560 | embryo sac egg cell differentiation | 0.00441 |
| GO:0010182 | sugar mediated signaling pathway | 0.00442 |
| GO:0051726 | regulation of cell cycle | 0.00446 |
| GO:1902275 | regulation of chromatin organization | 0.00504 |
| GO:0048440 | carpel development | 0.00623 |
| GO:0010072 | primary shoot apical meristem specificat... | 0.00652 |
| GO:0010162 | seed dormancy process | 0.00703 |

| | | |
|---|---|---|
| GO:0016567 | protein ubiquitination | 0.00725 |
| GO:0051604 | protein maturation | 0.00728 |
| GO:0051301 | cell division | 0.00770 |
| GO:0006281 | DNA repair | 0.00791 |
| GO:0010051 | xylem and phloem pattern formation | 0.00830 |
| GO:0048451 | petal formation | 0.00949 |
| GO:0048453 | sepal formation | 0.00949 |
| GO:0042127 | regulation of cell proliferation | 0.00961 |
| GO:0006497 | protein lipidation | 0.00973 |
| GO:0072528 | pyrimidine-containing compound biosynthe... | 0.00996 |
| GO:0000375 | RNA splicing, via transesterification re... | 0.01002 |
| GO:0000377 | RNA splicing, via transesterification re... | 0.01002 |
| GO:0006325 | chromatin organization | 0.01033 |
| GO:0006261 | DNA-dependent DNA replication | 0.01090 |
| GO:0031056 | regulation of histone modification | 0.01139 |
| GO:0051168 | nuclear export | 0.01186 |
| GO:0048229 | gametophyte development | 0.01219 |
| GO:0018377 | protein myristoylation | 0.01237 |
| GO:0031365 | N-terminal protein amino acid modificati... | 0.01237 |
| GO:0043603 | cellular amide metabolic process | 0.01280 |
| GO:2000242 | negative regulation of reproductive proc... | 0.01421 |
| GO:0000398 | mRNA splicing, via | 0.01444 |

| | | |
|---|---|---|
| | spliceosome | |
| GO:0019915 | lipid storage | 0.01513 |
| GO:0070646 | protein modification by small protein re... | 0.01522 |
| GO:0006498 | N-terminal protein lipidation | 0.01567 |
| GO:0006499 | N-terminal protein myristoylation | 0.01567 |
| GO:0006891 | intra-Golgi vesicle-mediated transport | 0.01834 |
| GO:0010224 | response to UV-B | 0.01854 |
| GO:0046519 | sphingoid metabolic process | 0.01864 |
| GO:0030048 | actin filament-based movement | 0.01866 |
| GO:0006928 | movement of cell or subcellular componen... | 0.01866 |
| GO:0010212 | response to ionizing radiation | 0.01882 |
| GO:0043604 | amide biosynthetic process | 0.01902 |
| GO:0070918 | production of small RNA involved in gene... | 0.01916 |
| GO:0031050 | dsRNA processing | 0.01916 |
| GO:0097435 | supramolecular fiber organization | 0.01928 |
| GO:0051302 | regulation of cell division | 0.01968 |
| GO:0006220 | pyrimidine nucleotide metabolic process | 0.01980 |
| GO:0006221 | pyrimidine nucleotide biosynthetic proce... | 0.01980 |
| GO:0009220 | pyrimidine ribonucleotide biosynthetic p... | 0.01980 |
| GO:0009218 | pyrimidine ribonucleotide metabolic proc... | 0.01980 |

| | | |
|---|---|---|
| GO:0016579 | protein deubiquitination | 0.01985 |
| GO:0009943 | adaxial/abaxial axis specification | 0.02030 |
| GO:0009944 | polarity specification of adaxial/abaxia... | 0.02030 |
| GO:0065001 | specification of axis polarity | 0.02030 |
| GO:0003002 | regionalization | 0.02159 |
| GO:0009553 | embryo sac development | 0.02206 |
| GO:0009955 | adaxial/abaxial pattern specification | 0.02224 |
| GO:0001510 | RNA methylation | 0.02231 |
| GO:0044275 | cellular carbohydrate catabolic process | 0.02232 |
| GO:0006518 | peptide metabolic process | 0.02252 |
| GO:0010048 | vernalization response | 0.02275 |
| GO:0000303 | response to superoxide | 0.02288 |
| GO:0000305 | response to oxygen radical | 0.02288 |
| GO:0045595 | regulation of cell differentiation | 0.02413 |
| GO:0009910 | negative regulation of flower developmen... | 0.02466 |
| GO:0032504 | multicellular organism reproduction | 0.02581 |
| GO:0000394 | RNA splicing, via endonucleolytic cleava... | 0.02586 |
| GO:0009057 | macromolecule catabolic process | 0.02587 |
| GO:0009311 | oligosaccharide metabolic process | 0.02601 |
| GO:0046520 | sphingoid | 0.02609 |

| | | |
|---|---|---|
| | biosynthetic process | |
| GO:0048439 | flower morphogenesis | 0.02634 |
| GO:0051338 | regulation of transferase activity | 0.02657 |
| GO:0033044 | regulation of chromosome organization | 0.02670 |
| GO:0043161 | proteasome-mediated ubiquitin-dependent ... | 0.02694 |
| GO:0010876 | lipid localization | 0.02704 |
| GO:0010351 | lithium ion transport | 0.02797 |
| GO:0051093 | negative regulation of developmental pro... | 0.02803 |
| GO:0006611 | protein export from nucleus | 0.02809 |
| GO:0048481 | plant ovule development | 0.02810 |
| GO:0035670 | plant-type ovary development | 0.02810 |
| GO:0001676 | long-chain fatty acid metabolic process | 0.02849 |
| GO:0005984 | disaccharide metabolic process | 0.02861 |
| GO:0009798 | axis specification | 0.02964 |
| GO:0043043 | peptide biosynthetic process | 0.02966 |
| GO:0006259 | DNA metabolic process | 0.02975 |
| GO:0006897 | endocytosis | 0.03171 |
| GO:0098657 | import into cell | 0.03171 |
| GO:0048193 | Golgi vesicle transport | 0.03182 |
| GO:0006412 | translation | 0.03248 |
| GO:0050826 | response to freezing | 0.03377 |
| GO:0009410 | response to xenobiotic stimulus | 0.03426 |
| GO:0007031 | peroxisome organization | 0.03434 |
| GO:0016571 | histone methylation | 0.03488 |

| | | |
|---|---|---|
| GO:0006508 | proteolysis | 0.03595 |
| GO:0050657 | nucleic acid transport | 0.03597 |
| GO:0050658 | RNA transport | 0.03597 |
| GO:0006403 | RNA localization | 0.03597 |
| GO:0006405 | RNA export from nucleus | 0.03597 |
| GO:0051236 | establishment of RNA localization | 0.03597 |
| GO:0065009 | regulation of molecular function | 0.03611 |
| GO:0000272 | polysaccharide catabolic process | 0.03805 |

Supplementary Table S6: Gene ontology enrichment analysis among genes with high additive variance $V_A$. The GOs ID, the term, and the p value of the significant GOs ($p < 0.03813$) are given.

| **GO.ID** | **Term** | ***P* value** |
|---|---|---|
| GO:0008150 | biological_process | 5.4e-12 |
| GO:0006007 | glucose catabolic process | 8.3e-06 |
| GO:0006096 | glycolytic process | 1.6e-05 |
| GO:0006094 | gluconeogenesis | 9.1e-05 |
| GO:0046686 | response to cadmium ion | 0.00015 |
| GO:0080129 | proteasome core complex assembly | 0.00016 |
| GO:0006511 | ubiquitin-dependent protein catabolic pr... | 0.00016 |
| GO:0051788 | response to misfolded protein | 0.00077 |
| GO:0009853 | photorespiration | 0.00099 |
| GO:0006412 | translation | 0.00231 |
| GO:0006626 | protein targeting to mitochondrion | 0.00243 |
| GO:0010498 | proteasomal protein catabolic process | 0.00382 |
| GO:0000162 | tryptophan biosynthetic process | 0.00545 |
| GO:0009805 | coumarin biosynthetic process | 0.00626 |
| GO:0006575 | cellular modified amino acid metabolic p... | 0.00627 |
| GO:0019760 | glucosinolate metabolic process | 0.00788 |
| GO:0042542 | response to hydrogen peroxide | 0.00846 |
| GO:0009651 | response to salt stress | 0.00865 |
| GO:0009610 | response to symbiotic fungus | 0.00903 |
| GO:0055114 | oxidation-reduction process | 0.00921 |
| GO:0019395 | fatty acid oxidation | 0.01062 |
| GO:1901566 | organonitrogen compound biosynthetic pro... | 0.01064 |
| GO:0034440 | lipid oxidation | 0.01098 |

| | | |
|---|---|---|
| GO:0042180 | cellular ketone metabolic process | 0.01139 |
| GO:0050801 | ion homeostasis | 0.01232 |
| GO:0006635 | fatty acid beta-oxidation | 0.01251 |
| GO:0009812 | flavonoid metabolic process | 0.01263 |
| GO:0007030 | Golgi organization | 0.01349 |
| GO:0010256 | endomembrane system organization | 0.01410 |
| GO:0000041 | transition metal ion transport | 0.01428 |
| GO:0030258 | lipid modification | 0.01529 |
| GO:0009408 | response to heat | 0.01625 |
| GO:0031365 | N-terminal protein amino acid modificati... | 0.01710 |
| GO:0006498 | N-terminal protein lipidation | 0.01733 |
| GO:0006499 | N-terminal protein myristoylation | 0.01733 |
| GO:0006520 | cellular amino acid metabolic process | 0.01776 |
| GO:0044272 | sulfur compound biosynthetic process | 0.01900 |
| GO:0072329 | monocarboxylic acid catabolic process | 0.01904 |
| GO:0048878 | chemical homeostasis | 0.01905 |
| GO:0006873 | cellular ion homeostasis | 0.01940 |
| GO:1901605 | alpha-amino acid metabolic process | 0.01967 |
| GO:1901420 | negative regulation of response to alcoh... | 0.01988 |
| GO:0009788 | negative regulation of abscisic acid-act... | 0.01988 |
| GO:1905958 | negative regulation of cellular response... | 0.01988 |
| GO:0016144 | S-glycoside biosynthetic process | 0.02042 |
| GO:0019761 | glucosinolate biosynthetic process | 0.02042 |
| GO:0019758 | glycosinolate biosynthetic process | 0.02042 |

| | | |
|---|---|---|
| GO:0046482 | para-aminobenzoic acid metabolic process | 0.02197 |
| GO:0022904 | respiratory electron transport chain | 0.02218 |
| GO:0009062 | fatty acid catabolic process | 0.02316 |
| GO:0030003 | cellular cation homeostasis | 0.02317 |
| GO:0000096 | sulfur amino acid metabolic process | 0.02332 |
| GO:0006497 | protein lipidation | 0.02407 |
| GO:0042157 | lipoprotein metabolic process | 0.02407 |
| GO:0042158 | lipoprotein biosynthetic process | 0.02407 |
| GO:0062012 | regulation of small molecule metabolic p... | 0.02437 |
| GO:0044281 | small molecule metabolic process | 0.02465 |
| GO:0044282 | small molecule catabolic process | 0.02651 |
| GO:0009813 | flavonoid biosynthetic process | 0.02669 |
| GO:0000097 | sulfur amino acid biosynthetic process | 0.02689 |
| GO:0006820 | anion transport | 0.02731 |
| GO:0055082 | cellular chemical homeostasis | 0.02735 |
| GO:0019752 | carboxylic acid metabolic process | 0.02824 |
| GO:0009060 | aerobic respiration | 0.02833 |
| GO:0006082 | organic acid metabolic process | 0.02871 |
| GO:0018377 | protein myristoylation | 0.02877 |
| GO:0046351 | disaccharide biosynthetic process | 0.02878 |
| GO:0044273 | sulfur compound catabolic process | 0.02948 |
| GO:0022613 | ribonucleoprotein complex biogenesis | 0.02958 |
| GO:0006833 | water transport | 0.03039 |
| GO:0042044 | fluid transport | 0.03039 |
| GO:0042773 | ATP synthesis | 0.03137 |

| | | |
|---|---|---|
| | coupled electron transport | |
| GO:0042775 | mitochondrial ATP synthesis coupled elec... | 0.03137 |
| GO:0043436 | oxoacid metabolic process | 0.03177 |
| GO:0006576 | cellular biogenic amine metabolic proces... | 0.03278 |
| GO:0042254 | ribosome biogenesis | 0.03338 |
| GO:0046395 | carboxylic acid catabolic process | 0.03374 |
| GO:0016054 | organic acid catabolic process | 0.03374 |
| GO:0055080 | cation homeostasis | 0.03403 |
| GO:1901659 | glycosyl compound biosynthetic process | 0.03423 |
| GO:0019321 | pentose metabolic process | 0.03462 |
| GO:0042732 | D-xylose metabolic process | 0.03462 |
| GO:0046149 | pigment catabolic process | 0.03562 |
| GO:0051596 | methylglyoxal catabolic process | 0.03602 |
| GO:0009438 | methylglyoxal metabolic process | 0.03602 |
| GO:0046185 | aldehyde catabolic process | 0.03602 |
| GO:0042182 | ketone catabolic process | 0.03602 |
| GO:0006661 | phosphatidylinositol biosynthetic proces... | 0.03637 |
| GO:0009963 | positive regulation of flavonoid biosynt... | 0.03637 |
| GO:0009269 | response to desiccation | 0.03800 |

**Supplementary Table S7: There was a core set of 2,829 genes that always clustered together in the k-clustering analysis.** Genes in this cluster had median $V_{NA}$ and $V_A$ values of 0.501 and 0.141, respectively. GO enrichment analysis highlights 294 GO terms (significance threshold *p* = 0.0077). The top processes are related to developmental functions, such as "gravitropism" (*p* = 6.4e-28) and "trichome morphogenesis" (*p* = 8.9e-15). Gene ontology enrichment analysis for the genes included in the cluster with high co-regulation. The GOs ID, the term, and the p value of the significant GOs (*p* < 0.077) are given.

| **GO.ID** | **Term** | ***P* value** |
|---|---|---|
| GO:0009630 | gravitropism | 6.4e-28 |
| GO:0010090 | trichome morphogenesis | 8.9e-15 |
| GO:0006486 | protein glycosylation | 9.7e-15 |
| GO:0006487 | protein N-linked glycosylation | 2.0e-14 |
| GO:0000956 | nuclear-transcribed mRNA catabolic proce... | 5.8e-14 |
| GO:0010228 | vegetative to reproductive phase transit... | 4.0e-12 |
| GO:0007155 | cell adhesion | 1.4e-11 |
| GO:0007062 | sister chromatid cohesion | 4.8e-11 |
| GO:0045010 | actin nucleation | 1.1e-10 |
| GO:0016926 | protein desumoylation | 2.4e-09 |
| GO:0016567 | protein ubiquitination | 2.5e-09 |
| GO:0016579 | protein deubiquitination | 2.5e-08 |
| GO:0050665 | hydrogen peroxide biosynthetic process | 2.8e-08 |
| GO:0048193 | Golgi vesicle transport | 3.8e-08 |
| GO:0010182 | sugar-mediated signaling pathway | 4.7e-08 |
| GO:0010072 | primary shoot apical meristem specificat... | 1.6e-07 |
| GO:0033044 | regulation of chromosome | 5.2e-07 |

| | | |
|---|---|---|
| | organization | |
| GO:0035196 | production of miRNAs involved in gene si... | 5.6e-07 |
| GO:0031048 | chromatin silencing by small RNA | 6.1e-07 |
| GO:0006897 | endocytosis | 6.3e-07 |
| GO:0000911 | cytokinesis by cell plate formation | 7.6e-07 |
| GO:0007010 | cytoskeleton organization | 8.6e-07 |
| GO:0009909 | regulation of flower development | 9.3e-07 |
| GO:0009887 | animal organ morphogenesis | 1.0e-06 |
| GO:0045132 | meiotic chromosome segregation | 1.0e-06 |
| GO:0045595 | regulation of cell differentiation | 1.1e-06 |
| GO:0009640 | photomorphogenesi s | 1.5e-06 |
| GO:0009845 | seed germination | 1.8e-06 |
| GO:0016558 | protein import into peroxisome matrix | 2.0e-06 |
| GO:0000278 | mitotic cell cycle | 2.3e-06 |
| GO:0000226 | microtubule cytoskeleton organization | 2.6e-06 |
| GO:0043687 | post-translational protein modification | 3.0e-06 |
| GO:0050826 | response to freezing | 6.3e-06 |
| GO:0008284 | positive regulation of cell proliferatio... | 6.7e-06 |
| GO:0010332 | response to gamma radiation | 6.8e-06 |
| GO:0010048 | vernalization response | 8.5e-06 |
| GO:0030244 | cellulose biosynthetic process | 1.9e-05 |
| GO:0009880 | embryonic pattern | 2.6e-05 |

| | | |
|---|---|---|
| | specification | |
| GO:0043247 | telomere maintenance in response to DNA ... | 2.9e-05 |
| GO:0006397 | mRNA processing | 2.9e-05 |
| GO:0019915 | lipid storage | 4.7e-05 |
| GO:0042138 | meiotic DNA double-strand break formatio... | 6.2e-05 |
| GO:0009616 | virus induced gene silencing | 6.4e-05 |
| GO:0008150 | biological_process | 6.5e-05 |
| GO:0032204 | regulation of telomere maintenance | 6.6e-05 |
| GO:0048765 | root hair cell differentiation | 6.7e-05 |
| GO:0010073 | meristem maintenance | 7.9e-05 |
| GO:0010267 | production of ta-siRNAs involved in RNA ... | 9.5e-05 |
| GO:0010162 | seed dormancy process | 0.00010 |
| GO:0007131 | reciprocal meiotic recombination | 0.00011 |
| GO:0010498 | proteasomal protein catabolic process | 0.00012 |
| GO:0051301 | cell division | 0.00012 |
| GO:0000398 | mRNA splicing, via spliceosome | 0.00013 |
| GO:0010050 | vegetative phase change | 0.00014 |
| GO:0009855 | determination of bilateral symmetry | 0.00018 |
| GO:0045893 | positive regulation of transcription, DN... | 0.00027 |
| GO:0000281 | mitotic cytokinesis | 0.00032 |
| GO:0009832 | plant-type cell wall biogenesis | 0.00038 |
| GO:0010638 | positive regulation | 0.00038 |

| | | |
|---|---|---|
| | of organelle organiz... | |
| GO:0006094 | gluconeogenesis | 0.00046 |
| GO:0006346 | methylation-dependent chromatin silencin... | 0.00048 |
| GO:0010051 | xylem and phloem pattern formation | 0.00048 |
| GO:0046855 | inositol phosphate dephosphorylation | 0.00053 |
| GO:0016571 | histone methylation | 0.00054 |
| GO:0048573 | photoperiodism, flowering | 0.00058 |
| GO:0006366 | transcription by RNA polymerase II | 0.00061 |
| GO:0051302 | regulation of cell division | 0.00061 |
| GO:0007033 | vacuole organization | 0.00061 |
| GO:0010014 | meristem initiation | 0.00066 |
| GO:0032957 | inositol trisphosphate metabolic process | 0.00068 |
| GO:0006499 | N-terminal protein myristoylation | 0.00074 |
| GO:0010315 | auxin efflux | 0.00080 |
| GO:0048825 | cotyledon development | 0.00081 |
| GO:0030422 | production of siRNA involved in RNA inte... | 0.00083 |
| GO:0051726 | regulation of cell cycle | 0.00098 |
| GO:0006468 | protein phosphorylation | 0.00100 |
| GO:0046777 | protein autophosphorylation | 0.00102 |
| GO:0006306 | DNA methylation | 0.00122 |
| GO:0009894 | regulation of catabolic process | 0.00139 |
| GO:0006342 | chromatin silencing | 0.00154 |

| | | |
|---|---|---|
| GO:0035194 | post-transcriptional gene silencing by RN... | 0.00173 |
| GO:0006635 | fatty acid beta-oxidation | 0.00177 |
| GO:0051273 | beta-glucan metabolic process | 0.00215 |
| GO:0007034 | vacuolar transport | 0.00224 |
| GO:0048364 | root development | 0.00260 |
| GO:0009793 | embryo development ending in seed dorman... | 0.00272 |
| GO:0090056 | regulation of chlorophyll metabolic proc... | 0.00275 |
| GO:0022904 | respiratory electron transport chain | 0.00275 |
| GO:0048437 | floral organ development | 0.00276 |
| GO:0009933 | meristem structural organization | 0.00302 |
| GO:0016192 | vesicle-mediated transport | 0.00310 |
| GO:0048229 | gametophyte development | 0.00342 |
| GO:0009910 | negative regulation of flower developmen... | 0.00354 |
| GO:0048580 | regulation of post-embryonic development | 0.00364 |
| GO:0090698 | post-embryonic plant morphogenesis | 0.00381 |
| GO:0016036 | cellular response to phosphate starvatio... | 0.00402 |
| GO:0006310 | DNA recombination | 0.00407 |
| GO:0007267 | cell-cell signaling | 0.00440 |
| GO:0006418 | tRNA aminoacylation for | 0.00468 |

| | | |
|---|---|---|
| | protein translat... | |
| GO:0016570 | histone modification | 0.00488 |
| GO:0010629 | negative regulation of gene expression | 0.00491 |
| GO:0042732 | D-xylose metabolic process | 0.00608 |
| GO:0009908 | flower development | 0.00635 |
| GO:0030865 | cortical cytoskeleton organization | 0.00637 |
| GO:0007129 | synapsis | 0.00663 |
| GO:0000394 | RNA splicing, via endonucleolytic cleava... | 0.00818 |
| GO:0048767 | root hair elongation | 0.00847 |
| GO:0016049 | cell growth | 0.00859 |
| GO:0040007 | growth | 0.00947 |
| GO:0032504 | multicellular organism reproduction | 0.00949 |
| GO:0010558 | negative regulation of macromolecule bio... | 0.01012 |
| GO:2000113 | negative regulation of cellular macromol... | 0.01012 |
| GO:0065007 | biological regulation | 0.01022 |
| GO:0042773 | ATP synthesis coupled electron transport | 0.01039 |
| GO:0042775 | mitochondrial ATP synthesis coupled elec... | 0.01039 |
| GO:0008356 | asymmetric cell division | 0.01039 |
| GO:0031329 | regulation of cellular catabolic process | 0.01039 |
| GO:0051641 | cellular localization | 0.01068 |
| GO:0010564 | regulation of cell | 0.01076 |

| | | |
|---|---|---|
| | cycle process | |
| GO:0031324 | negative regulation of cellular metaboli... | 0.01163 |
| GO:0006891 | intra-Golgi vesicle-mediated transport | 0.01205 |
| GO:0042546 | cell wall biogenesis | 0.01223 |
| GO:0009926 | auxin polar transport | 0.01313 |
| GO:0032879 | regulation of localization | 0.01334 |
| GO:0050789 | regulation of biological process | 0.01394 |
| GO:0070592 | cell wall polysaccharide biosynthetic pr... | 0.01400 |
| GO:0070589 | cellular component macromolecule biosynt... | 0.01400 |
| GO:0044038 | cell wall macromolecule biosynthetic pro... | 0.01400 |
| GO:0048827 | phyllome development | 0.01434 |
| GO:0006816 | calcium ion transport | 0.01438 |
| GO:0031327 | negative regulation of cellular biosynth... | 0.01442 |
| GO:0040008 | regulation of growth | 0.01452 |
| GO:0048439 | flower morphogenesis | 0.01493 |
| GO:0006312 | mitotic recombination | 0.01517 |
| GO:0098727 | maintenance of cell number | 0.01517 |
| GO:0019827 | stem cell population maintenance | 0.01517 |
| GO:0010413 | glucuronoxylan metabolic process | 0.01533 |
| GO:2000280 | regulation of root | 0.01585 |

| | | |
|---|---|---|
| | development | |
| GO:0009553 | embryo sac development | 0.01615 |
| GO:0045492 | xylan biosynthetic process | 0.01712 |
| GO:0031537 | regulation of anthocyanin metabolic proc... | 0.01719 |
| GO:0040029 | regulation of gene expression, epigeneti... | 0.01731 |
| GO:0044265 | cellular macromolecule catabolic process | 0.01746 |
| GO:0009890 | negative regulation of biosynthetic proc... | 0.01801 |
| GO:0016441 | posttranscriptional gene silencing | 0.01912 |
| GO:0097435 | supramolecular fiber organization | 0.01942 |
| GO:0051274 | beta-glucan biosynthetic process | 0.01965 |
| GO:0032875 | regulation of DNA endoreduplication | 0.02018 |
| GO:0090329 | regulation of DNA-dependent DNA replicat... | 0.02018 |
| GO:0006623 | protein targeting to vacuole | 0.02037 |
| GO:0072665 | protein localization to vacuole | 0.02037 |
| GO:0072666 | establishment of protein localization to... | 0.02037 |
| GO:0045491 | xylan metabolic process | 0.02119 |
| GO:0090567 | reproductive shoot system development | 0.02127 |
| GO:0051168 | nuclear export | 0.02146 |
| GO:0003002 | regionalization | 0.02164 |

| | | |
|---|---|---|
| GO:0030243 | cellulose metabolic process | 0.02194 |
| GO:0010359 | regulation of anion channel activity | 0.02233 |
| GO:0022898 | regulation of transmembrane transporter ... | 0.02233 |
| GO:0032412 | regulation of ion transmembrane transpor... | 0.02233 |
| GO:0032409 | regulation of transporter activity | 0.02233 |
| GO:0006119 | oxidative phosphorylation | 0.02293 |
| GO:0010074 | maintenance of meristem identity | 0.02300 |
| GO:0032880 | regulation of protein localization | 0.02300 |
| GO:0030029 | actin filament-based process | 0.02303 |
| GO:0010410 | hemicellulose metabolic process | 0.02360 |
| GO:0051172 | negative regulation of nitrogen compound... | 0.02522 |
| GO:1905393 | plant organ formation | 0.02551 |
| GO:0048449 | floral organ formation | 0.02583 |
| GO:0051640 | organelle localization | 0.02583 |
| GO:0006302 | double-strand break repair | 0.02587 |
| GO:0006007 | glucose catabolic process | 0.02587 |
| GO:0048509 | regulation of meristem development | 0.02598 |
| GO:0016246 | RNA interference | 0.02661 |
| GO:0061458 | reproductive system development | 0.02683 |
| GO:0048608 | reproductive | 0.02683 |

| | | |
|---|---|---|
| | structure development | |
| GO:0051240 | positive regulation of multicellular org... | 0.02794 |
| GO:0048522 | positive regulation of cellular process | 0.02990 |
| GO:0010540 | basipetal auxin transport | 0.03177 |
| GO:0045787 | positive regulation of cell cycle | 0.03177 |
| GO:1902532 | negative regulation of intracellular sig... | 0.03177 |
| GO:1902275 | regulation of chromatin organization | 0.03177 |
| GO:0090697 | post-embryonic plant organ morphogenesis | 0.03186 |
| GO:0048589 | developmental growth | 0.03326 |
| GO:0010383 | cell wall polysaccharide metabolic proce... | 0.03374 |
| GO:0019320 | hexose catabolic process | 0.03417 |
| GO:0046365 | monosaccharide catabolic process | 0.03417 |
| GO:0009749 | response to glucose | 0.03417 |
| GO:0090696 | post-embryonic plant organ development | 0.03424 |
| GO:0010304 | PSII associated light-harvesting complex... | 0.03490 |
| GO:0048582 | positive regulation of post-embryonic de... | 0.03495 |
| GO:0009314 | response to radiation | 0.03541 |
| GO:0052543 | callose deposition in cell wall | 0.03559 |

| | | |
|---|---|---|
| GO:2000243 | positive regulation of reproductive proc... | 0.03559 |
| GO:0031123 | RNA 3'-end processing | 0.03603 |
| GO:0009960 | endosperm development | 0.03603 |
| GO:0051253 | negative regulation of RNA metabolic pro... | 0.03779 |
| GO:0051049 | regulation of transport | 0.03794 |
| GO:0048638 | regulation of developmental growth | 0.03861 |
| GO:0006281 | DNA repair | 0.03991 |
| GO:0051094 | positive regulation of developmental pro... | 0.03994 |
| GO:0090693 | plant organ senescence | 0.04038 |
| GO:0010150 | leaf senescence | 0.04038 |
| GO:0045934 | negative regulation of nucleobase-contai... | 0.04114 |
| GO:0015931 | nucleobase-containing compound transport | 0.04220 |
| GO:0046467 | membrane lipid biosynthetic process | 0.04220 |
| GO:0006643 | membrane lipid metabolic process | 0.04233 |
| GO:0006874 | cellular calcium ion homeostasis | 0.04247 |
| GO:0048574 | long-day photoperiodism, flowering | 0.04247 |
| GO:0048440 | carpel development | 0.04253 |
| GO:0050657 | nucleic acid transport | 0.04370 |
| GO:0050658 | RNA transport | 0.04370 |
| GO:0006611 | protein export from | 0.04370 |

| | | |
|---|---|---|
| | nucleus | |
| GO:0006403 | RNA localization | 0.04370 |
| GO:0006405 | RNA export from nucleus | 0.04370 |
| GO:0051236 | establishment of RNA localization | 0.04370 |
| GO:0009416 | response to light stimulus | 0.04371 |
| GO:0052386 | cell wall thickening | 0.04384 |
| GO:0009225 | nucleotide-sugar metabolic process | 0.04384 |
| GO:0009057 | macromolecule catabolic process | 0.04399 |
| GO:0006338 | chromatin remodeling | 0.04431 |
| GO:1902531 | regulation of intracellular signal trans... | 0.04431 |
| GO:0010017 | red or far-red light signaling pathway | 0.04431 |
| GO:0071489 | cellular response to red or far red ligh... | 0.04431 |
| GO:0018205 | peptidyl-lysine modification | 0.04468 |
| GO:0050793 | regulation of developmental process | 0.04476 |
| GO:0007389 | pattern specification process | 0.04487 |
| GO:0048467 | gynoecium development | 0.04507 |
| GO:0009555 | pollen development | 0.04515 |
| GO:0071554 | cell wall organization or biogenesis | 0.04529 |
| GO:0010118 | stomatal movement | 0.04537 |
| GO:1903507 | negative regulation of nucleic acid-temp... | 0.04665 |
| GO:0045892 | negative regulation of transcription, DN... | 0.04665 |

| | | |
|---|---|---|
| GO:1902679 | negative regulation of RNA biosynthetic ... | 0.04665 |
| GO:0071555 | cell wall organization | 0.04707 |
| GO:0006974 | cellular response to DNA damage stimulus | 0.04809 |
| GO:0051604 | protein maturation | 0.05065 |
| GO:0043269 | regulation of ion transport | 0.05111 |
| GO:0034762 | regulation of transmembrane transport | 0.05145 |
| GO:0034765 | regulation of ion transmembrane transpor... | 0.05145 |
| GO:0007292 | female gamete generation | 0.05168 |
| GO:0050794 | regulation of cellular process | 0.05258 |
| GO:2000241 | regulation of reproductive process | 0.05267 |
| GO:0006511 | ubiquitin-dependent protein catabolic pr... | 0.05340 |
| GO:0009150 | purine ribonucleotide metabolic process | 0.05375 |
| GO:0055074 | calcium ion homeostasis | 0.05508 |
| GO:0002253 | activation of immune response | 0.05508 |
| GO:0048571 | long-day photoperiodism | 0.05508 |
| GO:0002218 | activation of innate immune response | 0.05508 |
| GO:0048518 | positive regulation of biological proces... | 0.05845 |
| GO:0042398 | cellular modified | 0.05945 |

| | | |
|---|---|---|
| | amino acid biosyntheti... | |
| GO:0007051 | spindle organization | 0.05945 |
| GO:0010608 | post-transcriptional regulation of gene e... | 0.05993 |
| GO:0021700 | developmental maturation | 0.06017 |
| GO:0010154 | fruit development | 0.06018 |
| GO:0044036 | cell wall macromolecule metabolic proces... | 0.06127 |
| GO:0019375 | galactolipid biosynthetic process | 0.06203 |
| GO:0043480 | pigment accumulation in tissues | 0.06203 |
| GO:0043481 | anthocyanin accumulation in tissues in r... | 0.06203 |
| GO:0043473 | pigmentation | 0.06203 |
| GO:0043476 | pigment accumulation | 0.06203 |
| GO:0043478 | pigment accumulation in response to UV l... | 0.06203 |
| GO:0043479 | pigment accumulation in tissues in respo... | 0.06203 |
| GO:0009560 | embryo sac egg cell differentiation | 0.06306 |
| GO:0000271 | polysaccharide biosynthetic process | 0.06317 |
| GO:0044070 | regulation of anion transport | 0.06375 |
| GO:1903959 | regulation of anion transmembrane transp... | 0.06375 |
| GO:0006406 | mRNA export from nucleus | 0.06376 |
| GO:0051028 | mRNA transport | 0.06376 |
| GO:0071427 | mRNA-containing | 0.06376 |

| | | |
|---|---|---|
| | ribonucleoprotein comple... | |
| GO:0043632 | modification-dependent macromolecule cat... | 0.06417 |
| GO:0019941 | modification-dependent protein catabolic... | 0.06417 |
| GO:0061647 | histone H3-K9 modification | 0.06542 |
| GO:0051567 | histone H3-K9 methylation | 0.06542 |
| GO:0009955 | adaxial/abaxial pattern specification | 0.06667 |
| GO:2001057 | reactive nitrogen species metabolic proc... | 0.06810 |
| GO:0031056 | regulation of histone modification | 0.06810 |
| GO:0010252 | auxin homeostasis | 0.06810 |
| GO:0016032 | viral process | 0.06810 |
| GO:0009954 | proximal/distal pattern formation | 0.06810 |
| GO:0010015 | root morphogenesis | 0.06830 |
| GO:0006275 | regulation of DNA replication | 0.06933 |
| GO:0072507 | divalent inorganic cation homeostasis | 0.06954 |
| GO:0016444 | somatic cell DNA recombination | 0.06963 |
| GO:0006473 | protein acetylation | 0.06963 |
| GO:0051645 | Golgi localization | 0.06963 |
| GO:0051646 | mitochondrion localization | 0.06963 |
| GO:0060151 | peroxisome localization | 0.06963 |
| GO:0019374 | galactolipid metabolic process | 0.06964 |
| GO:0009411 | response to UV | 0.06982 |
| GO:0043161 | proteasome- | 0.06990 |

| | | |
|---|---|---|
| | mediated ubiquitin-dependent ... | |
| GO:0051649 | establishment of localization in cell | 0.07344 |
| GO:0010646 | regulation of cell communication | 0.07411 |

a.

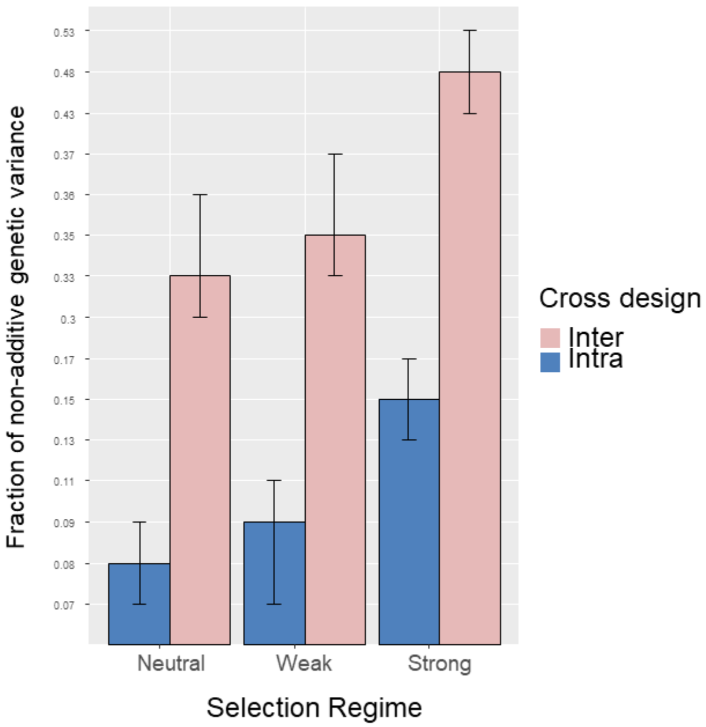

Fraction of non-additive genetic variance
0.53
0.48
0.43
0.37
0.36
0.35
0.33
0.3
0.17
0.15
0.13
0.11
0.09
0.08
0.07
Neutral
Weak
Strong
Selection Regime
Cross design
Inter
Intra

b.

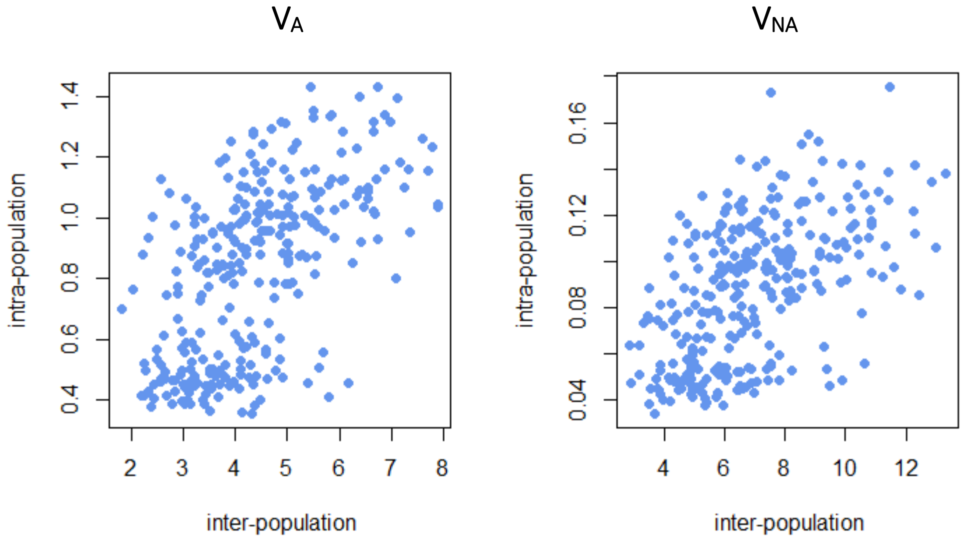

V_A
V_NA
intra-population
inter-population
0.4 0.6 0.8 1.0 1.2 1.4
2 3 4 5 6 7 8
0.04 0.08 0.12 0.16
4 6 8 10 12

c.

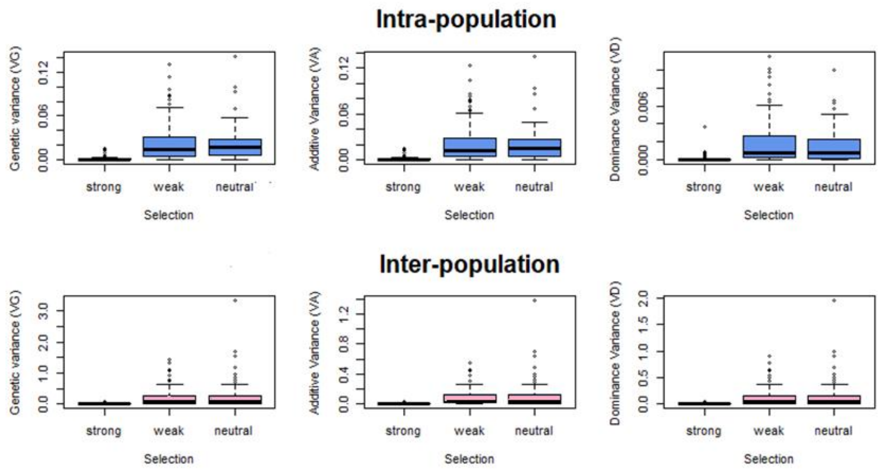

Intra-population
Inter-population
Genetic variance (VG)
Additive Variance (VA)
Dominance Variance (VD)
strong
weak
neutral
Selection

**Supplementary Figure S1:** Quantitative genetics model showing that non additive variance (modeled as dominance variance) is predicted to form a larger fraction of phenotypic variation in pedigrees obtained by crossing individuals from different populations and the action of selection can be better detected. Simulations assume population size of 200 and follow the model described in Clo & Opedal (2021) [38]. a. Summary of contribution of the non-additive variance to the total genetic variance ($V_{NA}/V_G$), as a function of the strength of purifying selection and the kind of progeny under study (mean ± 95% confidence interval computed over 100 repetitions - selection was modeled by modulating the stabilizing selection parameter $\omega^2$). Non-additive variance forms a larger fraction of the genetic variance in inter-population progenies, regardless of the strength of selection (see methods for details). This fraction also increases with the strength of selection, regardless of the cross type, but the effect of selection is stronger in inter-population crosses ($X^2 = 496.1$, df = 3, $p < 0.001$). b. Levels of additive and dominance variance expected in inter- vs. intrapopulation progenies are significantly correlated (Pearson correlation test: $V_A$ : $p < 10^{-16}$, $R^2 = 0.33$; $V_{NA} = V_D$ : $p < 10^{-16}$, $R^2 = 0.29$). c. Simulations of the amount of additive, dominance, and genetic variances when both gametes are sampled from the same population (intra-) vs when gametes are sampled each from a different population (inter-population progeny): non-additive variance forms a greater fraction of genetic variance when alleles were subjected to purifying selection.

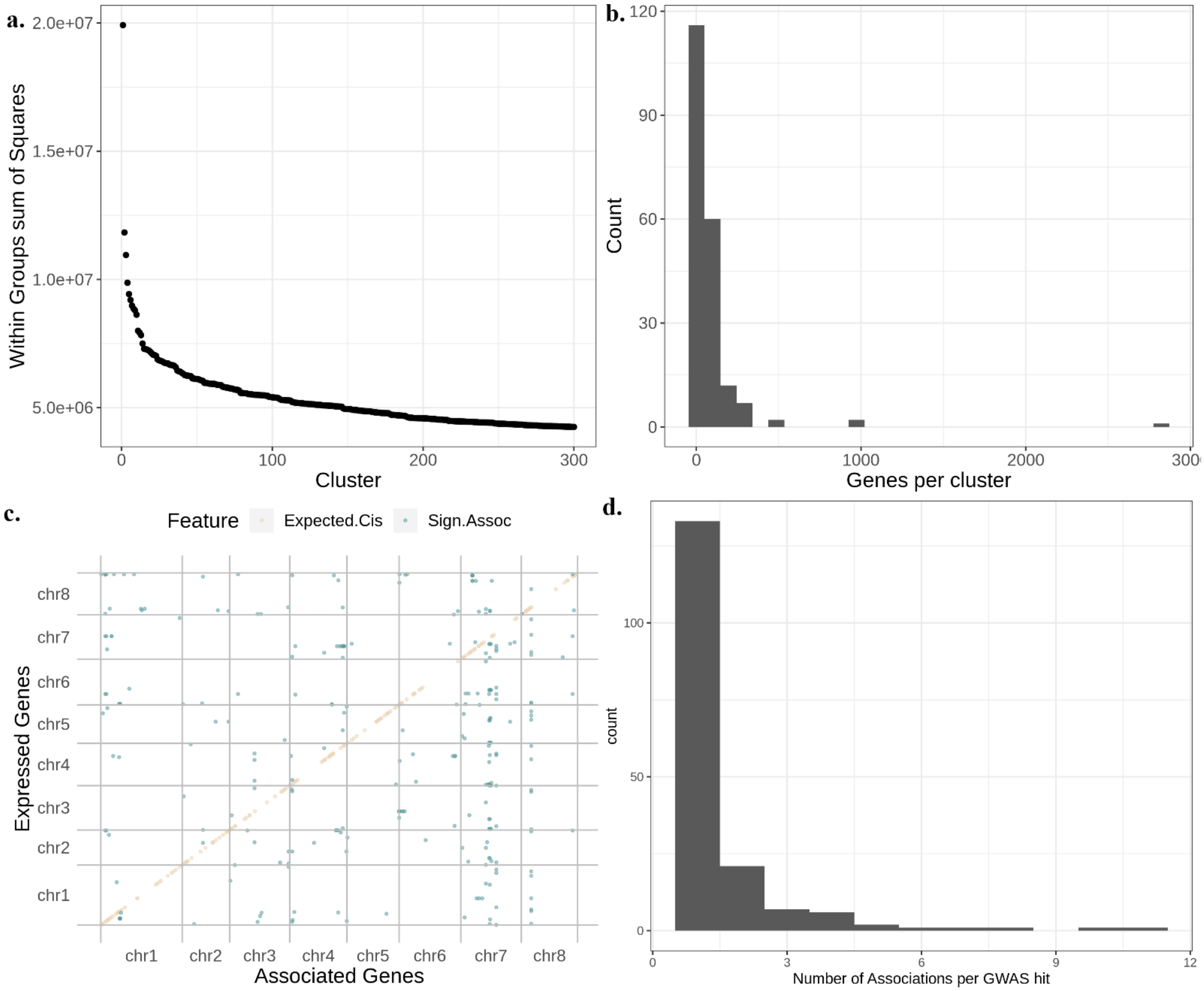

Supplementary Figure S2: Most genetic variance cannot be explained by large-effect variants. **a.** The within-groups sum of squares for each cluster partition was done based on the gene's co-expression. **b.** A histogram of the number of genes per cluster when we group them in 200 clusters based on their co-expression. **c.** The results of the GWAS run for each individual gene are summarized. On the y axis, the position of the expressed gene whose phenotype was tested is shown. On the x axis, the position along the chromosomes of the gene associating with the expression of at least one expressed gene is presented. Brown dots show the position of the expected association with oneself, blue dots show the significant associations of genetic variant with variation in transcript level. **d**. Histogram showing the number of transcripts associating with each variant.

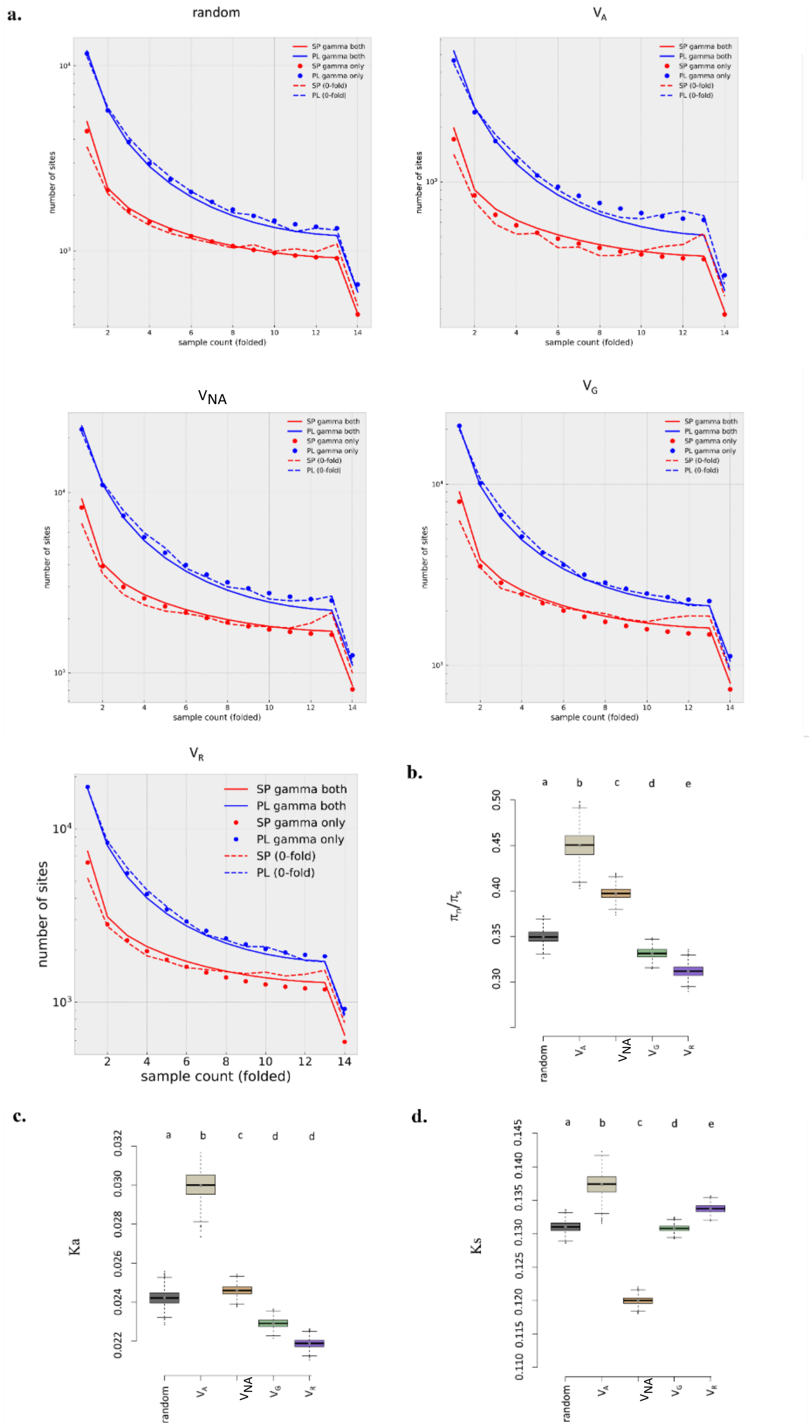
a.
random
VA
VNA
VG
VR
SP gamma both
PL gamma both
SP gamma only
PL gamma only
SP (0-fold)
PL (0-fold)
number of sites
sample count (folded)
b.
c.
d.
a b c d e
a b c d d
πn/πs
Ka
Ks
random
VA
VNA
VG
VR

**Supplementary Figure S3: Non-synonymous rates of evolution and their association with additive and non-additive variance in transcript expression. a.** The site-frequency spectrums for SP (red) and PL (blue) used to infer the DFE of the genes with high fraction of $V_A$, high fraction of $V_{NA}$, intermediate values of the fraction of $V_G$, high fraction of $V_R$, and a random set of genes shows a good fit to the expectations of the demographic model (line: observed, dots: predicted by the demographic model). **b.** We computed the mean and variance of average pi/ps values for each group based on population genomics datasets for SP. **c.** Divergence at non-synonymous sites, Ka, for genes grouped according to their genetic variance in transcript expression level. **d.** Divergence at synonymous sites, Ks, for genes grouped according to their genetic variance in transcript expression.

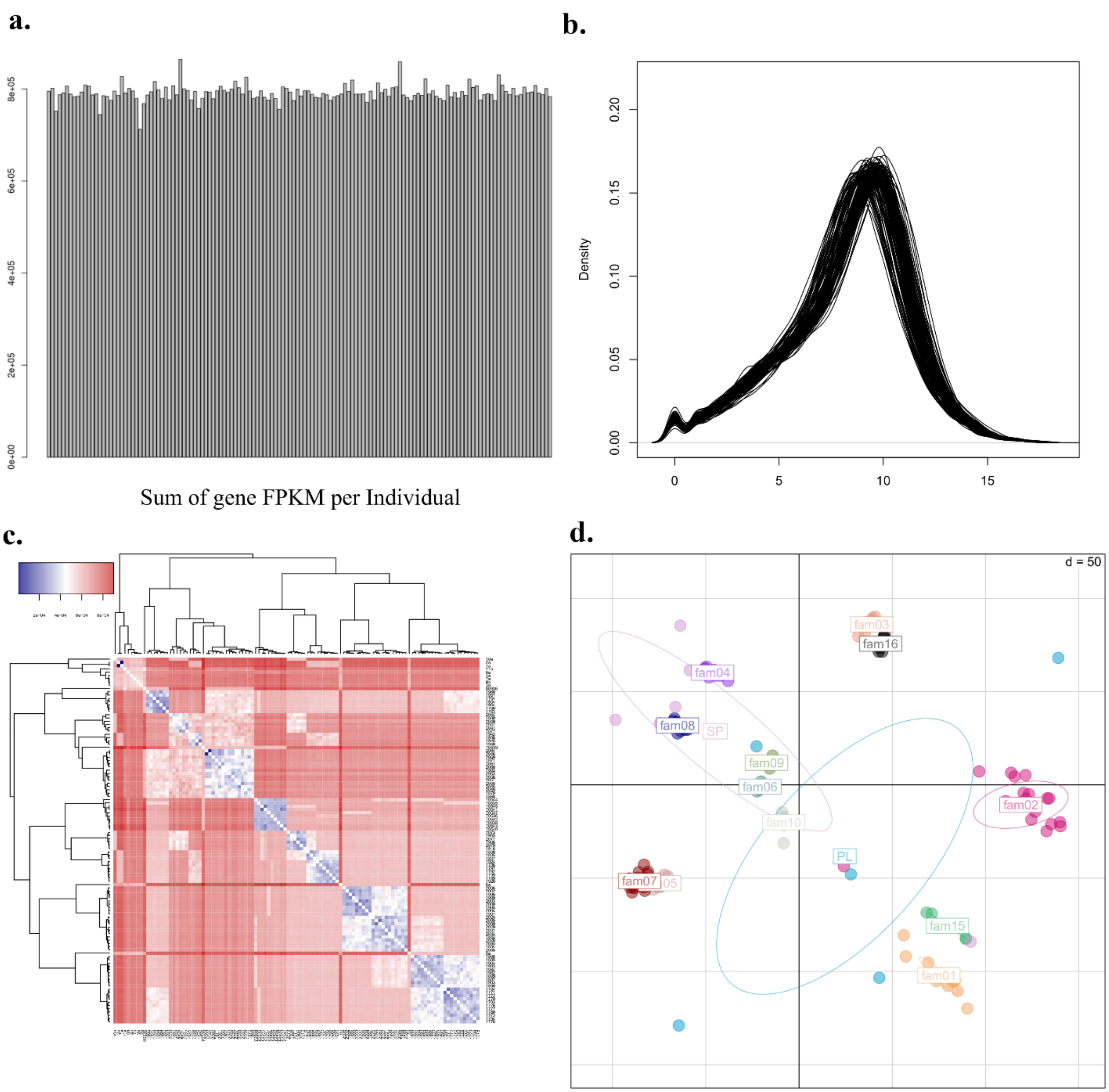

**Supplementary Figure S4: Quality control of the transcriptome samples.** **a.** The sum of the gene FPKM per sample is similar across all samples, which means that the gene counts between samples can be reliably compared. **b.** The log2 distribution of gene counts per sample is similar, indicating that there are no low-quality samples. **c.** The pairwise genetic distance of all samples was estimated. The parents' sequences are included in the analysis. **d.** Principal component analysis of the samples show that the full sibling individuals segregate together.
